# Identification and Engineering of UDP-rhamnosyltransferase from *Trillium tschonoskii* for Heterologous Biosynthesis of Polyphyllin II in Engineered Yeast

**DOI:** 10.1101/2025.01.26.634900

**Authors:** Yuxin Yang, Haowen Wang, Ziya Wu, Xing Wang, Yuru Tong, Wei Huang, Xuan Liu, Huan Zhao, Yating Hu, Xianan Zhang

**Author notes:** Authors for correspondence: Yating Hu Xianan Zhang.

## Abstract

Polyphyllins, a class of isospirostan-type steroidal saponins, exhibit potent cytotoxicity against a variety range of cancer cells. Although extensive research efforts have been made, the complete biosynthetic pathway of these compounds remains unclear. In order to elucidate these pathways, different tissues from Trillium tschonoskii were collected for sequencing that yielded 173,382 high-quality unigene sequences, among which 353 were annotated as glycosyltransferases. Then, a novel rhamnosyltransferase gene, UGT738A3, was characterized, which catalyzes the conversion of triglycoside polyphyllin III and pennogenin 3-O-beta-chacotrioside into tetraglycoside polyphyllin II and polyphyllin VII. The key residues that affect the catalytic activity of UGT738A3 were identified through site-directed mutation that the mutant A158T/P101L exhibited improved catalytic activity towards polyphyllin III and pennogenin 3-O-beta-chacotrioside by 2.5-fold and 6.5-fold respectively. We thus reconstructed the biosynthesis pathway of polyphyllin II in yeast by introducing the UGT93M3(catalyzing the formation of polyphyllin VI) and UGT738A3^A158T/P101L3^. This study not only elucidates the pivotal role of UGT738A3 in catalyzing the formation of tetraglycoside, but also provides highly efficient enzymatic components essential for the heterologous biosynthesis of polyphyllin saponins.

---

*Trillium tschonoskii*, a perennial herbaceous plant within the genus *Trillium* of the family Liliaceae, is employed in traditional medicine for its roots and rhizomes. Notably, it demonstrates significant efficacy in ameliorating central nervous system degenerative diseases, including Alzheimer’s disease(H. Y. Wang *et al*., 2024.). Its primary chemical components are steroidal saponins. In 1974, Nohara T(Kawasaki, 1975.) first isolated steroidal saponins from this herb. The main sapogenins identified include three types, diosgenin, pennogenin and furostanol-type sapogenins. The sapogenins typically form glycosidic bonds with various sugar moieties at the C3-OH position, including β-D-Glc, α-L-Rha, and α-L-Ara, resulting in triglycosides, tetraglycosides, and other oligosaccharides that exhibit diverse biological activities. Additionally, *T. tschonoskii* contains unique saponins such as trillenoside, kryptogenin saponin, and sterone *et al*(Yan *et al*., 2021.). Steroidal saponins possess intricate and diverse structures that exhibit significant structural similarities, rendering their chemical synthesis, isolation, and purification particularly challenging. Moreover, medicinal plants containing these compounds, such as those from the genus *Paris* and *Trillium*, are characterized by limited natural resources and extremely slow growth rates. For example, *T. tschonoskii* has been officially designated as a nationally protected rare and endangered plant in China(Q. Li *et al*., 2005.). This status poses significant challenges for the comprehensive research and development of polyphyllins. Currently, one promising strategy for achieving sustainable production of natural products is to engineer microorganisms by reconstructing the biosynthetic pathways of plant secondary metabolites within them(Tao *et al*., 2022.).

The biosynthetic pathway of diosgenin in plants has been thoroughly analyzed. Analogous to terpenoids, the biosynthesis of steroidal saponins involves upstream genes from both the MEP and MVA pathways, with the MVA pathway being predominant. 2,3-Oxidosqualene serves as a pivotal branch point in both the steroid and terpenoid metabolic pathways. In the steroid biosynthetic pathway, cycloartenol synthase (CAS) catalyzes the cyclization of 2,3-oxidosqualene to generate cycloartenol, which is a critical precursor for the biosynthesis of steroid compounds. Cycloartenol was further modified by oxidation and reduction to produce cholesterol, the key precursor of diosgenin(Salisbury *et al*., 2023.). Yin(Yin *et al*., 2018.) identified *Pp*CYP90B27, which catalyzes the hydroxylation of cholesterol at the C-22 position to form 22R-hydroxycholesterol. Subsequently, *Pp*CYP90G4 continue to catalyze the hydroxylation at the C-16 position of cholesterol, and 16S,22R-dihydroxycholesterol undergoes cyclization and dehydrogenation to form the E-ring(Christ *et al*., 2019, Zhou *et al*., 2021). Following this, C-26 hydroxylation by *Pp*CYP94D108 and spontaneously cyclization lead to generation of diosgenin(Christ *et al*., 2019). Despite the fact that penogenin merely contains an additional C-17 hydroxyl group compared to diosgenin, the key enzyme responsible for catalyzing this transformation has yet to be identified (Fig. 1).

**Fig. 1.**
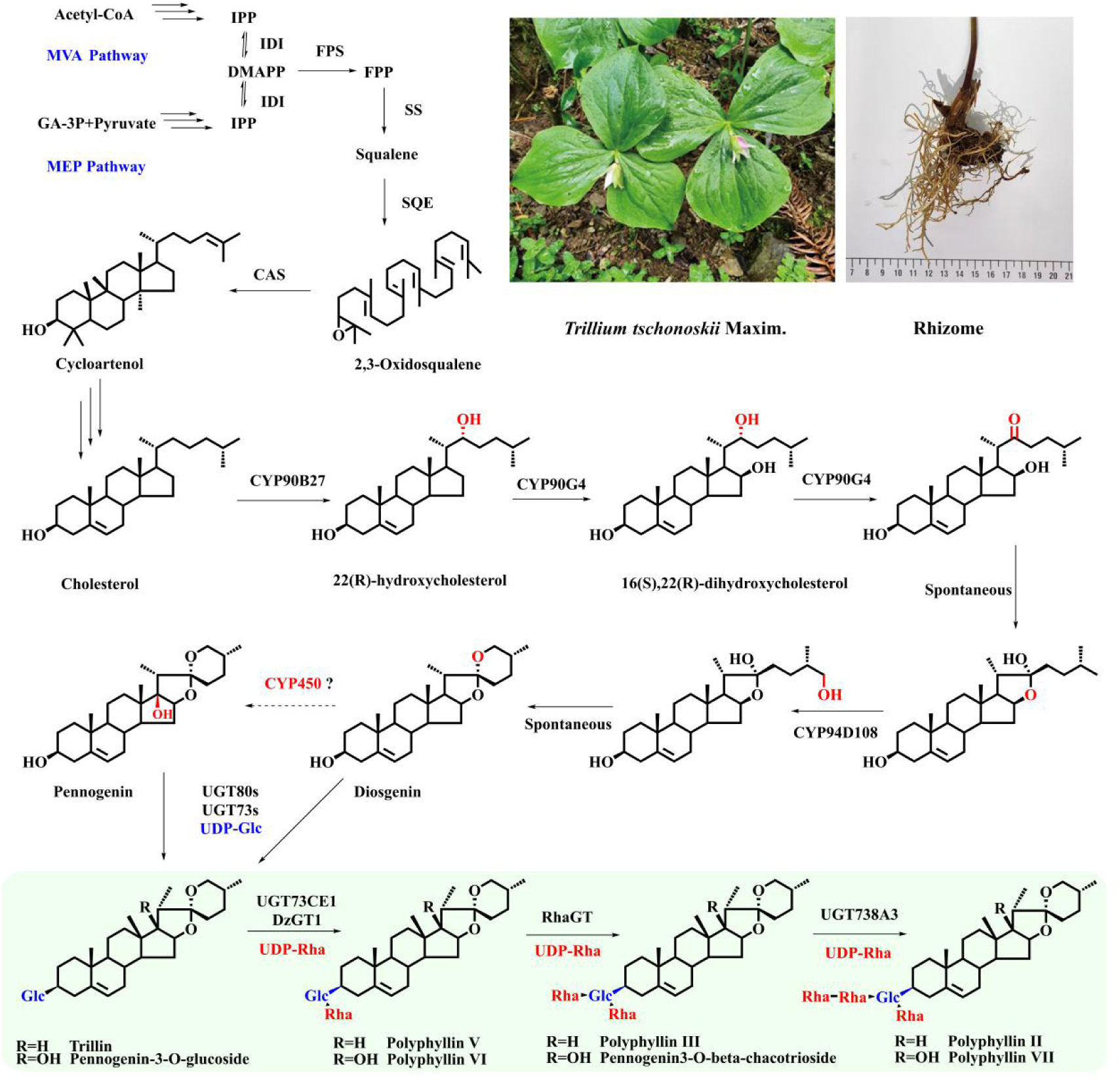
The biosynthetic pathways of polyphyllin.

In previous studies, several glycosyltransferases (GTs) responsible for the glucosylation of the C3-OH group of steroid sapogenins have been identified. These enzymes catalyze the C3 hydroxyl glucosylation of diosgenin and pennogenin, resulting in the formation of trillin and penogenin-3-O-glucoside(Song, *et al*.,2022, Chen, *et al*., 2023, He *et al*., 2023.). Subsequently, a 2’-O-rhamnosyltransferase UGT73CE1(Chen *et al*., 2023.) and DzGT1(Li *et al*., 2021), was identified to catalyze the addition of a rhamnose moiety at the 2’-O position, thereby generating polyphyllin V and polyphyllin VI(Fig.1). Nevertheless, to date, the number of reported sugar-sugar rhamnosyltransferases(RhaGTs) remains limited(Brandt *et al*., 2021.), which restricts the efficient production of polyphyllins through microbial metabolic engineering.

In this study, we conducted a comprehensive analysis of the transcriptome data from various tissues of *T. tschonoskii* to identify and confirm that UGT738A3 catalyzes the conversion of polyphyllin III and pennogenin 3-O-beta-chacotrioside(PRRG) into tetraglycoside polyphyllins II and VII. Through homologous sequence alignment, molecular docking, and mutant construction, we successfully enhanced the enzymatic activity of UGT738A3 *in vitro*. Using molecular dynamics(MD) simulations and docking studies, we elucidated the mechanism underlying the increased activity of the A158T/P101L mutant. Additionally, our findings indicate that the substrate specificity of the UGT738A3 A158T/P101L mutant was significantly broadened. By integrating UGT738A3 with a GT identified in the biosynthetic pathway of steroidal glycoalkaloids, we demonstrated that polyphyllin V can be effectively converted to polyphyllin II in *Saccharomyces cerevisiae*.

## Materials and Methods

### Plant materials

The plant material of *T. tschonoskii* was collected in April 2023 from Baoxing County, Ya’an City, Sichuan Province. All plant samples were immediately frozen in liquid nitrogen after collection and stored at -80°C. Each sample had three independent biological replicates for sequencing.

### Transcriptome sequencing of *T. tschonoskii*

Novogene Co., Ltd. was commissioned to perform RNA sequencing of *T. tschonoskii* using the Illumina platform. Raw sequencing data contained a small proportion of reads with adapters or low-quality sequences. To obtain clean reads, we filtered out adapter-containing reads, reads with ambiguous bases (N), and low-quality reads (where more than 50% of bases had a Qphred score ≤ 20). High-quality reads were then assembled into transcripts, and the longest unigenes were selected as reference sequences. The Corset program was used for hierarchical clustering of the transcripts, and the clustered sequences served as references for further analysis. Gene function annotation was conducted using seven databases: NR, NT, KO, SwissProt, PFAM, GO, and KOG.

### Determination of metabolite contents in distinct tissues of *T. tschonoskii*

Five distinct tissues of *T. tschonoskii* were retrieved from a -80°C freezer and dried to constant weight at 40°C. The dried samples were subsequently ground into fine powder and extracted with methanol to achieve a final concentration of 50 mg/mL. After ultrasonic treatment for 2 hours, the mixture was centrifuged, and the supernatant was filtered obtain the sample solution. Quantitative analysis of polyphyllin III, protodioscin, polyphyllin VII, and PRRG was performed using an ultra-performance liquid chromatography (UPLC) system (U3000). The chromatographic conditions were as follows: a Waters BEH-C18 reversed-phase column (1.7 μm, 2.1 mm × 100 mm) was used with water (A) and acetonitrile (B) as the mobile phase. The elution gradient was programmed as follows: 0–8 min, 40%–47% B; 8–15 min, 47%–80% B; 15–17 min, 80%–100% B; 17–18 min, 100% B; 18–19 min, 100%–40% B; 19–24 min, 40% B. The flow rate was maintained at 0.3 mL/min, with an injection volume of 5 μL. The column temperature was kept constant at 40°C. All solvents were procured from Beijing Honghu United Chemical Products Co., Ltd., while the analytical standards were sourced from Shanghai Yuanye BioTechnology Co., Ltd (Shanghai, China).

### Functional identification of UGT738A3

The substrates for the enzymatic reactions included polyphyllin V, polyphyllin III, polyphyllin VI, and PRRG, with UDP-Rha serving as the glycosyl donor. Diosgenin and pennogenin were used as positive controls with UDP-Glu as the glycosyl donor, while a blank sample served as the negative control. The *in vitro* enzymatic reactions were conducted for 16 hours. Catalytic products were analyzed using LC-MS (SYNAPT G2-Si). Chromatographic conditions were as follows: a Waters BEH-C18 reversed-phase column (1.7 μm, 2.1 mm × 100 mm) was used with water (A) and acetonitrile (B) as the mobile phase. The elution gradient was programmed as: 0–9 min, 30%–60% B; 9–11 min, 60% B; 11–13 min, 60%–70% B; 13–16 min, 70%–100% B; 16–18 min, 100% B; 18–18.5 min, 100%–30% B; 18.5–23 min, 30% B. The flow rate was at 0.3 mL/min, with an injection volume of 5 μL. The column temperature was maintained at 30°C. Full-MS analysis was performed in positive ion mode with a scanning range of 100–1000 Da and a scan time of 0.2 s.

### Expression and purification of UGT738A3

The full-length UGT738A gene was cloned into the pET28a(+) vector, incorporating an N-terminal 6xHis tag, and subsequently transformed into *E. coli* BL21(DE3) for expression. Single colonies were cultured in 40 mL LB medium supplemented with 50 μg/mL kanamycin at 37°C with shaking at 250 rpm overnight. The culture was then diluted 1:100 into 4 L of LB medium containing 50 μg/mL kanamycin and grown until OD_600_ reached 0.6∼0.8. Protein expression was induced with 1 mM IPTG, followed by incubation for 20 hours at 16°C with shaking at 250 rpm. Following cell disruption, the supernatant was collected and loaded onto a Ni-NTA column, Protein purification was performed using a linear gradient of imidazole (10–500 mM). The purified protein was subsequently concentrated using a 30 kDa ultrafiltration centrifugal device and stored at -80°C.

### The influence of pH, temperature and time

Due to the enhanced catalytic activity of the mutant protein, its optimal reaction conditions were investigated with respect to pH, temperature, and reaction time. Enzymatic reactions were conducted for 3 hours at various temperatures and in different buffer systems to evaluate the optimal conditions. The pH values tested included: 4.0–6.0 using sodium citrate buffer, 6.0–8.0 using phosphate buffer, and 7.0–9.0 using Tris-HCl buffer. Reactions were performed at temperatures of 25°C, 30°C, 37°C, 40°C, 45°C, and 50°C. Additionally, the effect of reaction time wasevaluated by conductingreactions at eight different time points: 0.5, 1, 2, 4, 6, 8, 16, and 20 hours. The enzymatic reaction mixture consisted of 1 μL UDPG (100 mM), 2 μL polyphyllin III (5 mM), 5 μg of purified protein, and buffer to a final volume of 50 μL. After the reaction, 100 μL of ice-cold methanol was added to terminate the reaction.

### Enzyme kinetics research

Enzyme kinetics assays were conducted in a final reaction volume of 50 μL, containing 50 mM Tris-HCl (pH 8.0), 5 μg of purified enzyme, 2 mM UDP-Rha, and varying concentrations of polyphyllin III and PRRG(raning from 1.25 to 320 μM). Reactions were incubated at 30°C for 4 hours, after which 100 μL of ice-cold methanol was added to terminate the reaction. The mixture was then centrifuged at 14,000 rpm for 20 minutes, and the supernatant was analyzed by LC-MS/MS using an AB SCIEX 6500 system. All experiments were performed in triplicate. Michaelis-Menten plots were generated to analyze the kinetic data and determine the kinetic parameters.

### Substrate promiscuity and sugar donor heterogeneity

Each reaction was conducted in a final volume of 200 μL, containing 50 mM Tris-HCl (pH 8.0), 188 μL of crude enzyme extract, 0.1 mM sugar donor, and 0.2 mM substrate. The reactions were incubated at 30°C for 16 hours and terminated by adding 600 μL of ice-cold methanol. The mixture were then centrifuged at 14,000 rpm for 20 minutes, and the supernatants were collected and analyzed by LC-MS using a SYNAPT G2-Si system.

### Homology Modeling, Molecular Docking and Site-Directed Mutagenesis of UGT738A3

The three-dimensional structure of UGT738A3 was modeled using AlphaFold 3. fSubsequently, molecular docking simulations were performed with Autodock Vina to predict the binding interactions between the enzyme and its substrates. The docking results were visualized and analyzed using PyMOL software. Mutation primers were synthesized by Tianyi Huiyuan Biotechnology Co., Ltd. The mutant was expressed and purified following the procedures described above, and its catalytic activity was verified through functional assays. The samples were analyzed by LC-MS/MS using an AB SCIEX 6500 system.

### MD simulation

MD simulations were conducted using the Gromacs 2018.4 software package under constant temperature and pressure conditions, employing periodic boundary conditions. The Amber14SB all-atom force field and TIP3P water model were utilized for the simulations. During the MD simulation, all bonds involving hydrogen atoms were constrained using the LINCS algorithm, allowing an integration time step of 2 fs. Long-range electrostatic interactions were calculated using the Particle-mesh Ewald (PME) method. Non-bonded interactions were truncated at 10 Å, with a neighbor list update every 10 steps. The system temperature was maintained at 298.15 K using the V-rescale thermostat, and pressure was controlled at 1 bar using the Parrinello-Rahman barostat. Initially, energy minimization was performed on the systems using the steepest descent method to eliminate unfavorable atomic contacts. Subsequently, 1 ns equilibration simulations were carried out in the NVT and NPT ensembles at 298.15 K. Finally, a 300 ns production MD simulation was performed, with snapshots saved every 10 ps. The simulation results were analyzed using using Gromacs tools and visualized with VMD.

### Functional verification in yeast cells

The genes encoding UGT738A3 and UGT93M3 were introduced into an engineered yeast straincapable of expressing polyphyllin V. The transformed yeast cells were plated on Ura dropout plates and cultured for three days. After confirming plasmid maintenance, the yeast cells were inoculated into 100 ml YPD medium at an initial OD_600_ of 0.05 and cultured for five days at 30°C with shaking at 200 rpm. The cells harvested were then lysed using a high-pressure homogenizer at 1200 psi. The lysate was subjected to two sequential extractions with an equal volume of n-butanol. The conbined organic extracts were concentrated to a final volume of 1 ml. The prepared samples were analyzed using an AB SCIEX 6500 system to assess the enzymatic activity and product formation.

## Results

### Transcriptome sequencing, functional annotation and RhaGT genes screening

A total of 15 sequencing libraries were generated using Illumina sequencing, encompassing three biological replicates from five distinct plant tissues, rhizome, fibrous root, stem, leaf, and flower. The clean reads were assembled using Trinity, resulting in 173,382 high-quality unigene sequences. The N50 length of the assembled unigenes was 1005 bp (Fig. S1a,b). To obtain comprehensive gene function information, annotations were performed using seven databases. The results showed that all 173,382 high-quality unigene sequences were annotated. A total of 117,039 (67.5%) sequences were annotated in at least one database, while 17,077 (9.84%) sequences were annotated in all seven databases (Fig. S1c). After performing KO annotation, genes were classified based on their involvement in specific KEGG metabolic pathways (Fig. S1d). Specifically, 723 sequences were associated with terpenoid and polyketide metabolism, which may related to the biosynthesis of steroidal saponins in *T. tschonoskii*.

By analyzing the transcriptome annotation data, we filtered out genes with FPKM expression levels less than 5 and sequence lengths shorter than 1200 bp, resulting in the identification of 353 GT genes. A phylogenetic tree was constructed by incorporating these genes with previously reported RhaGTs from *P.polyphylla*, *D. polystachya*, and *G. uralensis* (Fig. 2b, S3; protein sequences in Table S1). A total of 60 candidate RhaGTs were screened out, which mainly belong to the UGT73clan, UGT89clan, and UGT91clan.

**Fig. 2.**
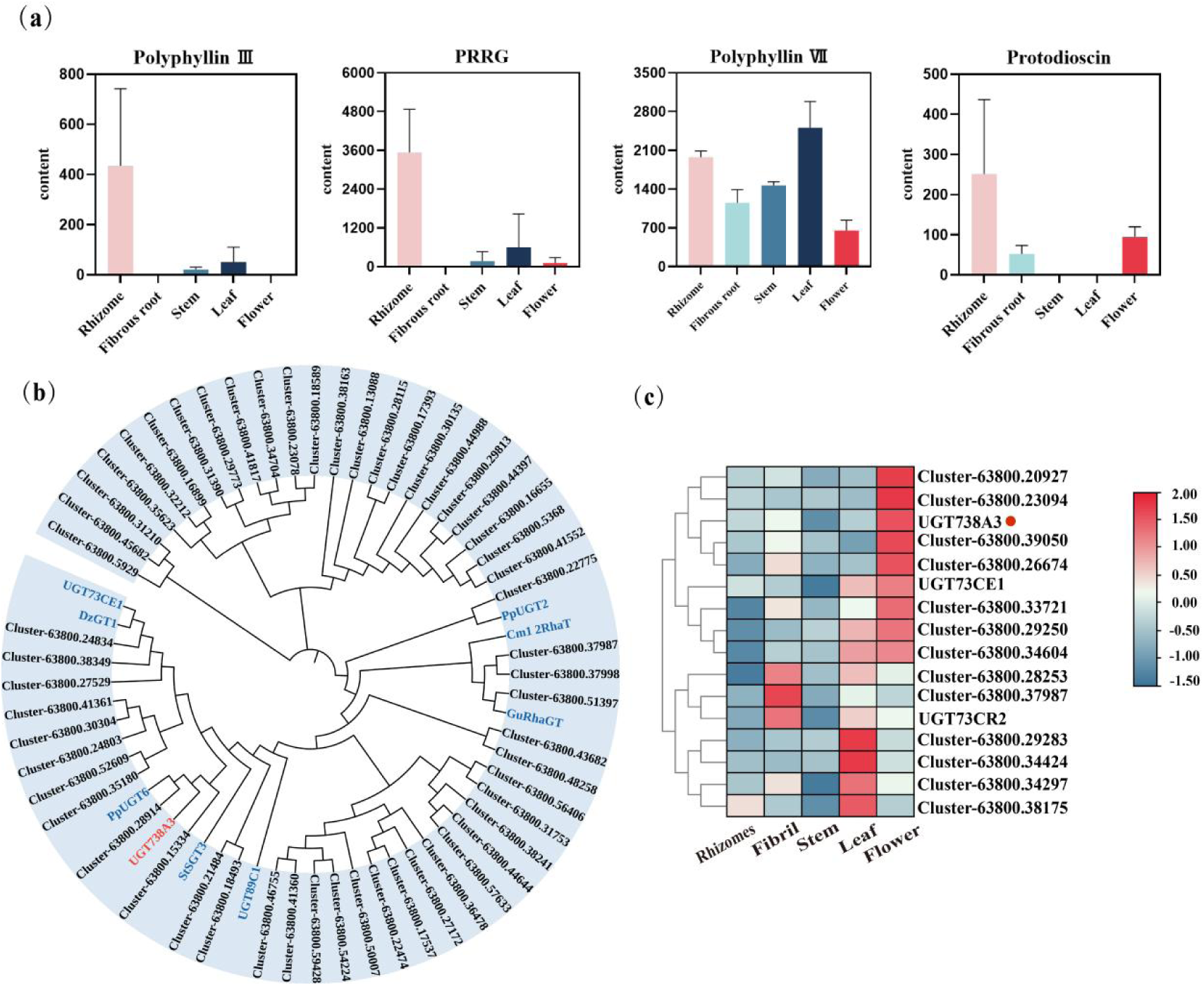
Screening of RhaGTs based on the metabolite content, gene annotation and expression levels in distanct tissues of *T. tschonoskii*. (a) Determination of polyphyllin III, protodioscin, polyphyllin VII and PRRG in five distanct parts of *T. tschonoskii.* (b) The phylogenetic placement of UGT738A3 within the phylogenetic tree. (c) The expression profile of UGT738A3 as depicted in the heatmap.

Analysis of the polyphyllin content across distanct parts of *T. tschonoskii*(Fig. 2a) revealed that, compared to triglucosides, the tetrasaccharide compounds polyphyllin VII and protodioscin exhibited higher levels in the flowers, while the accumulation of polyphyllin III and PRRG was most pronounced in the rhizome. Based on the heatmap depicting the expression levels of 60 candidate RhaGT genes(Fig. 2c, S4), functional verification will be prioritized for those genes with higher expression levels in flowers.

### Functional Characterization of the RhaGT Genes

Using mixed cDNA from the stems, leaves, and flowers of *T. tschonoski*, we successfully cloned and identified a RhaGT gene *Tt*UGT39, which was officially named UGT738A3 by the International Glycosyltransferase Nomenclature Committee. The ORF of UGT738A3 is 1488 bp, encoding a protein of 495 amino acids with a molecular weight of 54 kDa. When fused with an maltodextrin binding protein (MBP) tag, the protein’s molecular weight is approximately 100 kDa(Fig. S5b). Through *in vitro* enzymatic reactions and comparison with standards, it was found that UGT738A3 can convert polyphyllin III and PRRG into polyphyllin II and polyphyllin VII, respectively (Fig. 3a, b, c). Enzymatic property analysis demonstrated that the enzyme exhibited maximum conversion rate at 30°C in a Na_2_HPO_4_-NaH_2_PO_4_ buffer solution with a pH of 7.0 after a reaction time of 16h. Kinetic analysis showed that the enzyme has a higher affinity for polyphyllin III, as evidenced by the lower Km value (Table 1, Fig. S6).

**Fig. 3.**
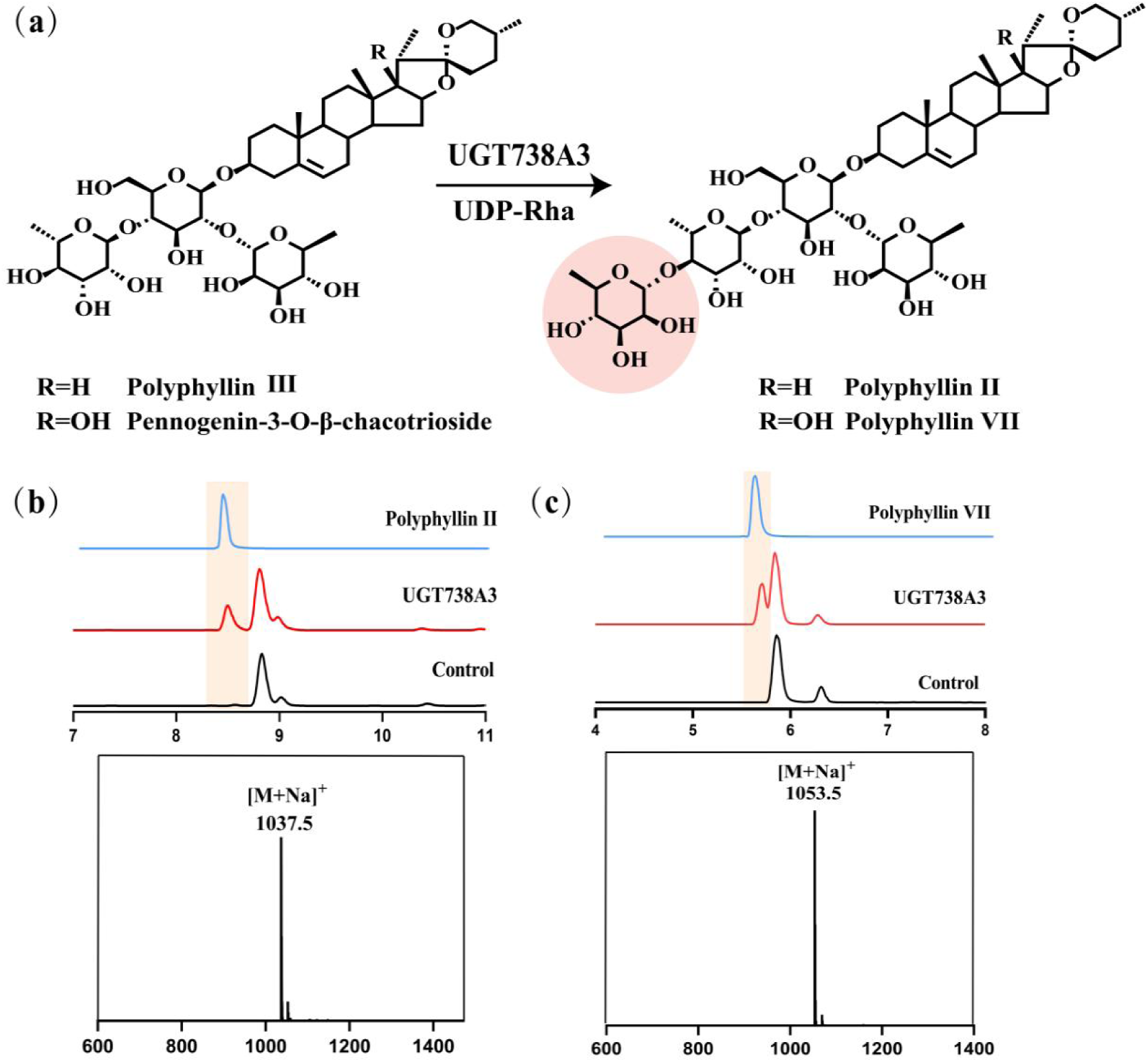
Functional Characterization of UGT738A3 from *T. tschonoski* . (a) UGT738A3 catalyzes the conversion of polyphyllin III and PRRG into polyphyllin II and polyphyllin VII. (b) HPLC and mass spectrometry analysis confirming UGT738A3’s role in converting polyphyllin III to polyphyllin II. (c) HPLC and mass spectrometry analysis validating UGT738A3’s function in transforming PRRG into polyphyllin VII.

**Table 1.** Kinetic Parameters of UGT738A3 and Its Mutants.

| Name | Substrate | K<sub>m</sub>(μM) | V<sub>max</sub>(nM/min) | K<sub>cat</sub>(s<sup>-1</sup>) |
| --- | --- | --- | --- | --- |
| UGT738A3 | Polyphyllin III | 9.26 | 0.09 | 0.008 |
| UGT738A3 | PRRG | 203.3 | 0.23 | 0.02 |
| UGT738A3<sup>A158T/P101L</sup> | Polyphyllin III | 14.32 | 8.78 | 1.62 |
| UGT738A3<sup>A158T/P101L</sup> | PRRG | 17.21 | 15.1 | 2.80 |

To investigate the substrate promiscuity and sugar donor selectivity of UGT738A3, a comprehensive set of five sugar donors and fourteen substrates (as detailed in Fig. S8) was selected for analysis. UPLC-MS/MS analysis revealed no catalytic reactions could occur between UGT738A3 and other sugar donors or substrates. These results indicate that UGT738A3 exhibits high regioselectivity specifically towards the 4’’-OH group and demonstrates high substrate specificity for UDP-Rha.

### Molecular mechanism of catalytic activity of UGT738A3

To elucidate the molecular mechanism underlying the catalytic activity of UGT738A3, we initially compared its structure with previously reported three-dimensional crystal structures using SWISS-MODEL. This analysis revealed a maximum similarity of only 34.87%. Attempts to determine its crystal structure were unsuccessful due to inadequate yields of monomeric protein. Consequently, we utilized AlphaFold 3 for protein modeling and conducted docking analysis using Autodock Vina, employing UDP-Rha as the sugar donor and polyphyllin III as the substrate (Fig. 4a). Currently, only a limited number of crystal structures of RhaGTs have been reported. Notably, *Arabidopsis* UGT89C1 represents the first RhaGT for which a crystal structure has been elucidated. Studies indicate that His22 and Asp121 are highly conserved residues that may constitute a receptor-His-Asp complex during catalytic process. Pro147, Ile148, Asp356, and His357 play a critical role in determining the specificity of UGT89C1 toward rhamnose (Zong *et al*., 2019.). Our docking results indicate that Gln356 and Ala357 in UGT738A3 form hydrogen bonds with the hydroxyl group of UDP-Rha. Mutagenesis experiments revealed that substituting these residues with alanine led to a substantial loss of catalytic function, thereby confirming the essential role of Gln356 and Ala357 in the catalytic activity of UGT738A3 (Fig. 4b).

**Fig. 4.**
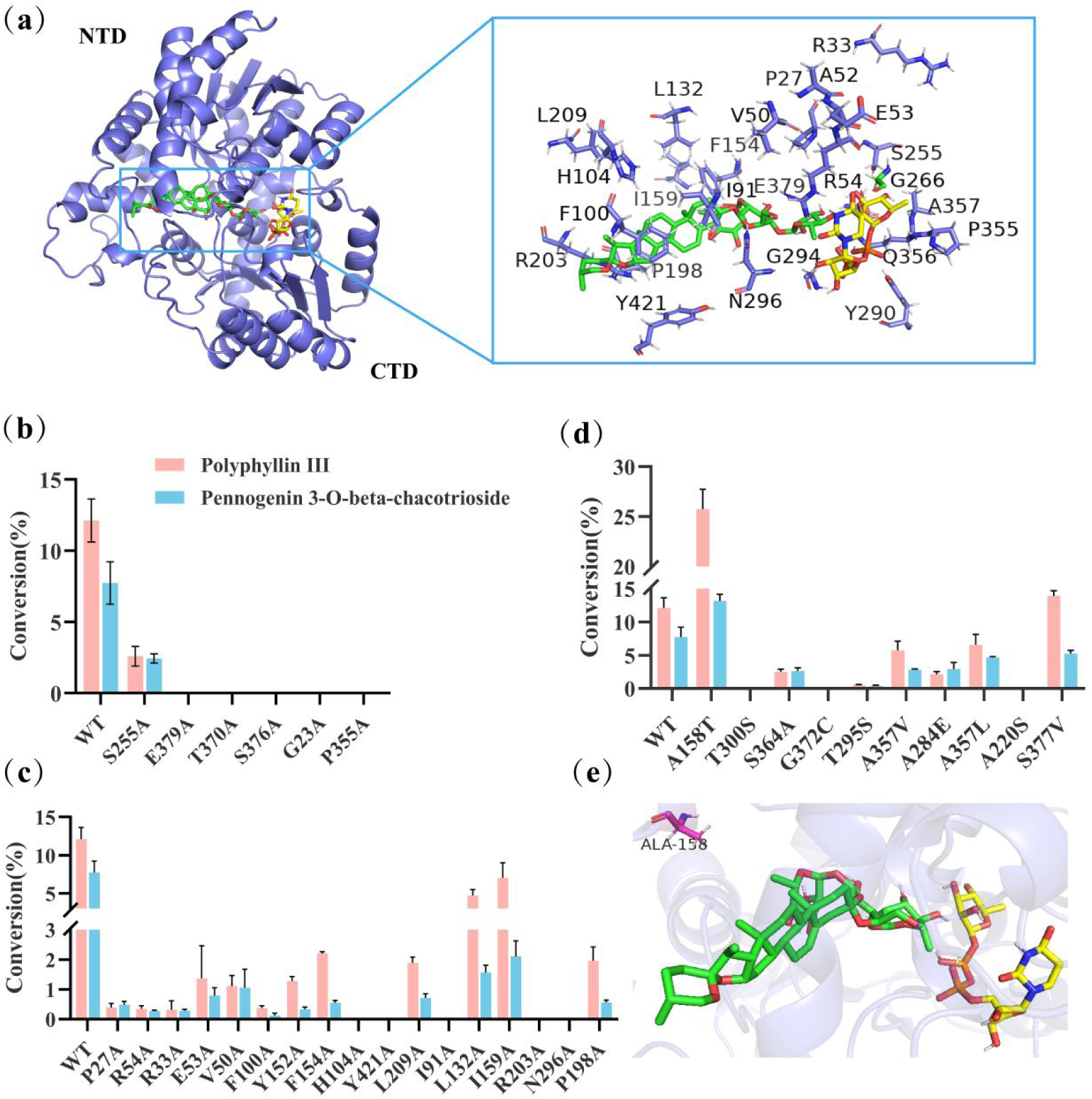
The results of protein modeling, molecular docking and site-directed mutagenesis of UGT738A3. (a) Structural modeling of UGT738A3 protein and molecular docking analysis with polyphyllin III and UDP-Rha. (b) Based on the results of molecular docking, alanine scanning was conducted on the amino acids near UDP-Rha. (c) Perform alanine scanning on the residues located in 5 Å of the substrate channel for polyphyllin III. (d) Mutual substitution of different amino acids in homologous sequence alignment. (e) The position of A158 residue in molecular docking.

### Identify the key residues that influence the catalytic activity of UGT738A3

The conversion efficiencies of UGT738A3 in catalyzing the formation of polyphyllin II and polyphyllin VII from their respective substrates are relatively low, at 12.11% and 7.73% respectively, thereby presenting presenting a significant challenge for large-scale production of these compounds. Consequently, we implemented enzyme engineering modifications on UGT738A3, guided by sequence differences and molecular docking results, to enhance its catalytic activity. Since the substrate must traverse the substrate channel to enter the binding pocket, the dimensions and volume of the tunnel entrance are crucial determinants of the enzyme’s catalytic activity(G. Y. Li *et al*., 2017.). Based on the molecular docking results, we identified key amino acids in a 5Å radius of the substrate channel for alanine scanning (Fig. 4c). The analysis revealed that most of the mutants exhibited diminished activity, with mutations at positions H104, Y421, I91, R203, and N296 resulting in complete loss of function. Notably, residues R203 and N296 form hydrogen bonds with the substrate polyphyllin III, indicating their critical role in stabilizing the substrate conformation (Fig. 4a).

The UGT738A3 sequence was aligned with previously reported RhaGTs(McCue *et al*., 2007; J. Li *et al*., 2021; Z. L. Wang *et al*., 2022; Xue *et al*., 2024.)(Table S1, Fig. S9), and the differing amino acids were reciprocally substituted(Fig. 4d). The results demonstrated that mutant A158T exhibited a twofold increase in the conversion rate of polyphyllin III compared to the wild type and an 80% increase in the conversion rate of PRRG. In contrast, the mutant S337V exhibited only a marginal improvement in the conversion rate of polyphyllin III (Fig. 4d). Subsequently, saturation mutagenesis of A158T and double mutation of A158T/S377V were performed to identify superior variants (Fig. S10), however, none surpassed the performance of the A158T mutant. Additionally, when prosapogenin B was used as the substrate and UDP-Rha as the sugar donor, the A158T mutant generated a peak with a molecular weight of 891.4 at 10.38 min (Fig. S11b, c). This peak did not match the retention time of standard polyphyllin III, suggesting that the potential site for sugar addition in the product may be the 4’’-hydroxyl group. Molecular docking analysis indicated that in the A158T mutant, the distance between the binding sites of prosapogenin B and UDP-Rha was reduced to 2.8 Å, compared to 16.2 Å in the wild type(Fig. S11d, e). This substantial reduction in distance likely accounts for the diminished catalytic activity observed in the wild type.

To elucidate the mechanism underlying the enhanced activity of the A158T mutant, molecular docking simulations were conducted following structural modeling. The results indicated that the A158 residue is positioned relatively distant from the substrate (Fig. 4e), suggesting that it may enhance catalytic efficiency by stabilizing the protein structure. To test this hypothesis, a 300-ns MD simulation was performed.

The RMSD (Root Mean Square Deviation) values of all protein atoms stabilized within 50 ns, with fluctuations consistently remaining below 1 Å. This indicates a stable and reliable system throughout the simulation period (Fig. 5a). The radius of gyration (Rg) value of the A158T mutant protein exhibited minimal fluctuation over the entire simulation duration, suggesting highly stable protein conformation (Fig. 5b). In contrast, the WT protein displayed more pronounce fluctuations, indicating comparatively lower stability relative to the A158T mutant (Fig. 5b). The RMSF values futher corroborated the enhanced stability of the A158T protein structure (Fig. 5c). Moreover, the calculated binding free energy for A158T was -64.9064 kcal/mol, which is notably lower than that of WT (-56.8695 kcal/mol), indicating a more stable complex formation(Fig. 5e). These results collectively indicate that the A158T mutant exhibits superior catalytic activity, attributed to its stable protein conformation and favorable substrate binding configuration.

**Fig. 5.**
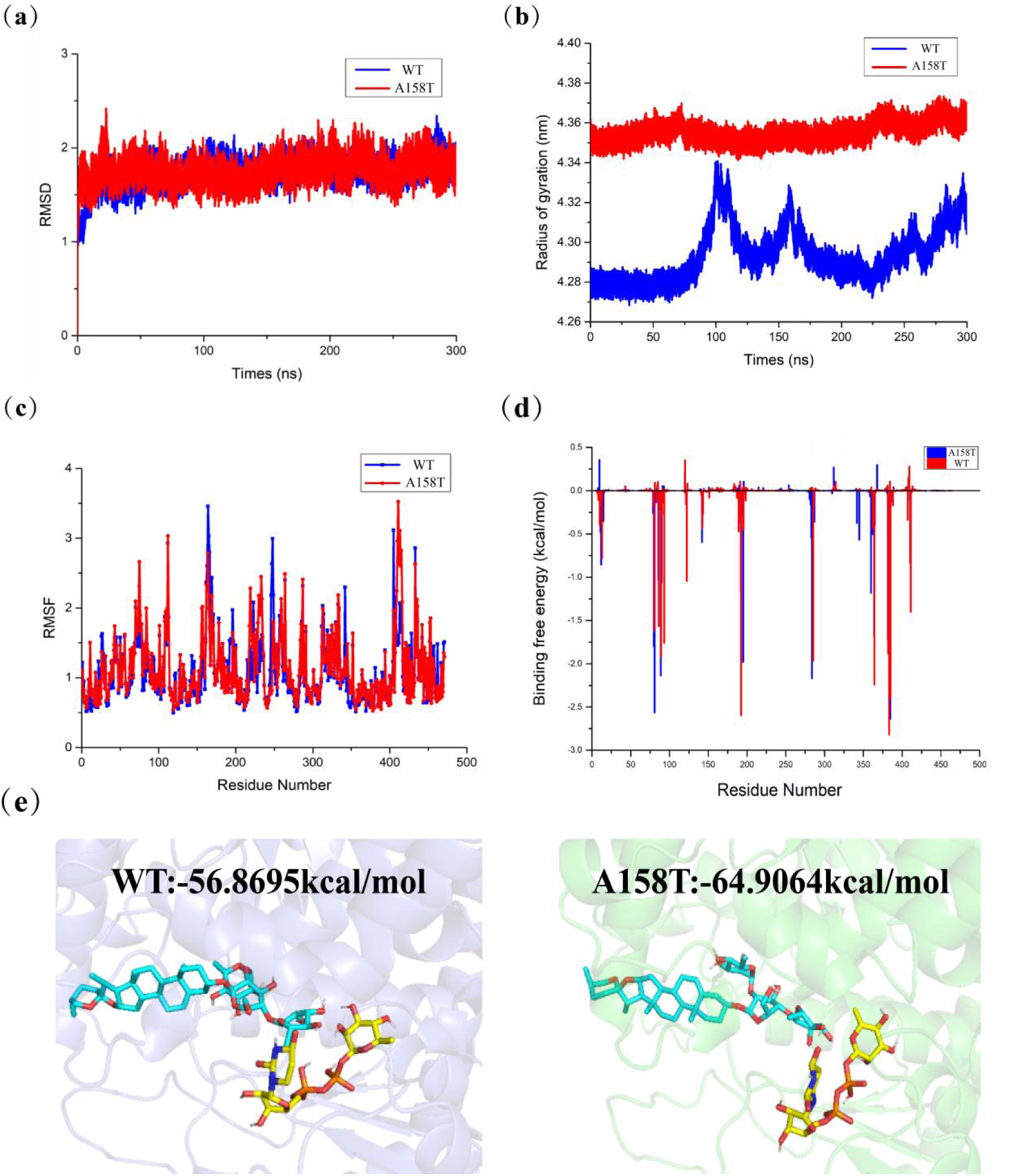
Molecular dynamics simulation results of A158T mutant and wild-type proteins in complex with polyphyllin III revealed significant differences in the structural stability. (a) The RMSD of protein molecules within the complex system as a function of simulation time. (b) The variation in Rg of the composite system over the course of the simulation. (c) RMSF analysis of amino acid residues in protein complexes. (d) Contribution of each amino acid to the binding free energy in the complex. (e) Docking outcomes and binding free energy calculations for both WT and A158T mutant systems.

### Focusing on rational iterative site-specific mutations (FRISM)

The FRISM process is an iterative methodology that involves the systematic screening of a limited number of predicted mutant enzymes to enhance their selectivity and catalytic activity, the highest-performing mutant from each iteration becomes the starting point for the next cycle of optimization(D. Y. Li *et al*., 2020.). Previously, we identified the A158T mutant with enhanced catalytic activity and utilized it as the foundation for further optimization. Based on docking simulations, we targeted amino acids in proximity to the substrate channel for additional mutations. Given that the MBP tag may influence protein activity, we reconstructed the His-pET28a-UGT738A3 and His-pET28a-UGT738A3^A158T^ recombinant proteins without the MBP tag (Fig. S7). The results showed that removal of the MBP tag significantly increased the activity of the A158T mutant towards polyphyllin III by approximately twofold(Fig. 6a). In the second round of saturation mutagenesis, mutating the P101 site to Ala, Leu, or Phe resulted in enhanced enzyme activity. Notably, the P101L mutant achieved conversion rates of 73.15% for polyphyllin III and 44.75% for PRRG, with kinetic data indicating superior catalytic efficiency (Table 1). Based on molecular docking results, we conducted a third round of mutagenesis on the A158T/P101L background. However, none of the resulting mutants exhibited higher activity than UGT738A3^A158T/P101L^.

**Fig. 6.**
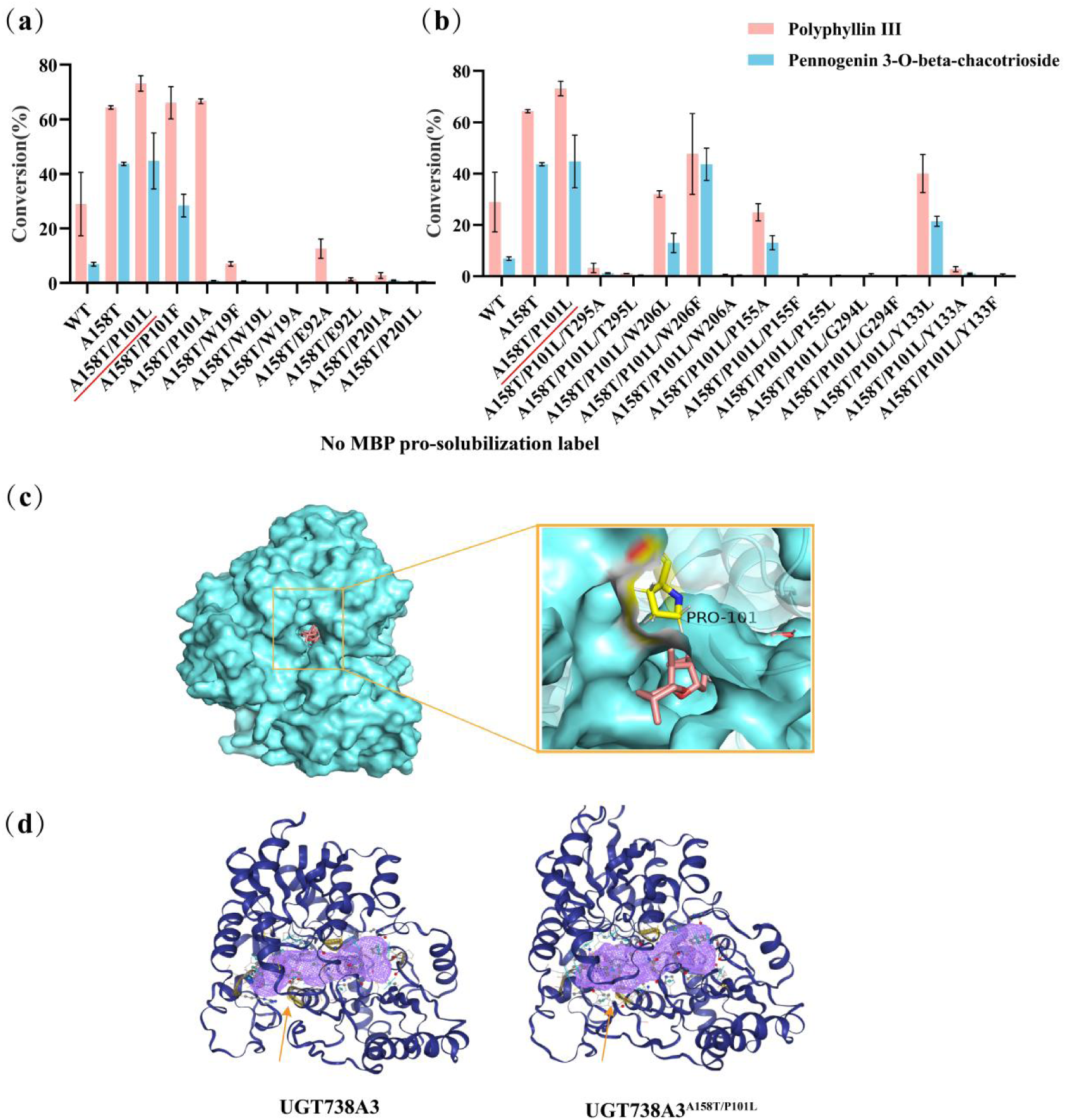
Iterative mutations based on the A158T background, guided by the FRISM strategy and informed by molecular docking results. (a) The double mutation results of UGT738A3 based on A158T. (b) The triple mutation outcomes of UGT738A3 based on A158T/P101L. (c) The position of P101 residue in molecular docking. (d) Pocket volume predictions for UGT738A3 and UGT738A3^A158T/P101L^ mutant.

The UGT738A3^A158T/P101L^ protein was modeled using AlphaFold 3 and subsequently docked with polyphyllin III. Analysis revealed that the P101 residue is positioned at the entrance of the substrate channel (Fig. 6c). Crystal structure analysis of UGT74AN3 indicates that Phe is a highly rigid residue, resulting in a narrow binding pocket and a near-complete loss of catalytic activity when mutated to Phe(Huang *et al*., 2023.). Therefore, mutating P101 to Ala or Leu decreases the side chain size, whereas mutating to Phe enhances side chain flexibility. This increases the channel pocket volume and facilitates substrate entry, thereby enhancing catalytic activity. Pocket prediction analysis using Proteins.Plus (https://proteins.plus) (Table. S2, Fig. 6d) revealed that both WT and mutant proteins exhibit a pocket depth of 25.62 Å. However, the pocket volume increased from 612.86 Å³ in the WT to 740.86 Å³ in the mutant. This increase in pocket volume facilitates substrate entry and significantly enhances catalytic activity. Previously reported GT pockets typically range from 800 to 1500 Å³(Brazier-Hicks *et al*., 2007; Modolo *et al*., 2009; T *et al*., 2015; KM *et al*., 2016; Thompson *et al*., 2017; Maharjan *et al*., 2020; Graef *et al*., 2023.), whereas the smaller pocket of UGT738A3 restricts substrate entry and impairs catalysis. Using the A158T/P101L mutant, further substrate spectrum analysis of the compounds shown in Fig. S8 revealed a prominent peak at 1217.4 m/z in the mass spectrum when protodioscin was used as the substrate. This suggests that the product may undergo glycosylation at the 4’’-hydroxy position, resulting in the formation of dichotomin(Fig. S12a,b).

### The complete biosynthesis of polyphyllin II in yeast

In 2024, Prashant D. Sonawane’s team identified the RhaGT genes UGT93M2 and UGT93M3 while investigating the biosynthetic pathway of steroid glycoside alkaloids(Lucier *et al*., 2024.). Considering the structural similarity between C3-OH sugar chain of β1-solamargine and that of polyphyllin V, the functional identification of UGT93M3 was conducted using polyphyllin V and polyphyllin VI as substrates. Through UPLC-MS/MS and comparison with the standards, it was found that UGT93M3 catalyzes the glycosylation of polyphyllin V and polyphyllin VI to produce polyphyllin III and PRRG (Fig. 7b,c). Subsequently, UGT93M3 was used as a probe to screen candidate genes in the *T. tschonoski* transcriptome database (Fig. S13a), and the candidate genes were cloned and functionally identified. Unfortunately, none of these genes had the same function as UGT93M3. We transformed the UGT738A3^A158T/P101L^ mutant and UGT93M3(TZ42) into the a genetically engineered yeast strain capable of producing polyphyllin V stored in our lab(Fig. 7a). Following extraction and analysis of the products, the results were consistent with those of the *in vitro* reactions. By comparison with the standard, polyphyllin II was detected at a retention time of 5.35 minutes(Fig. 7d,e).

**Fig. 7.**
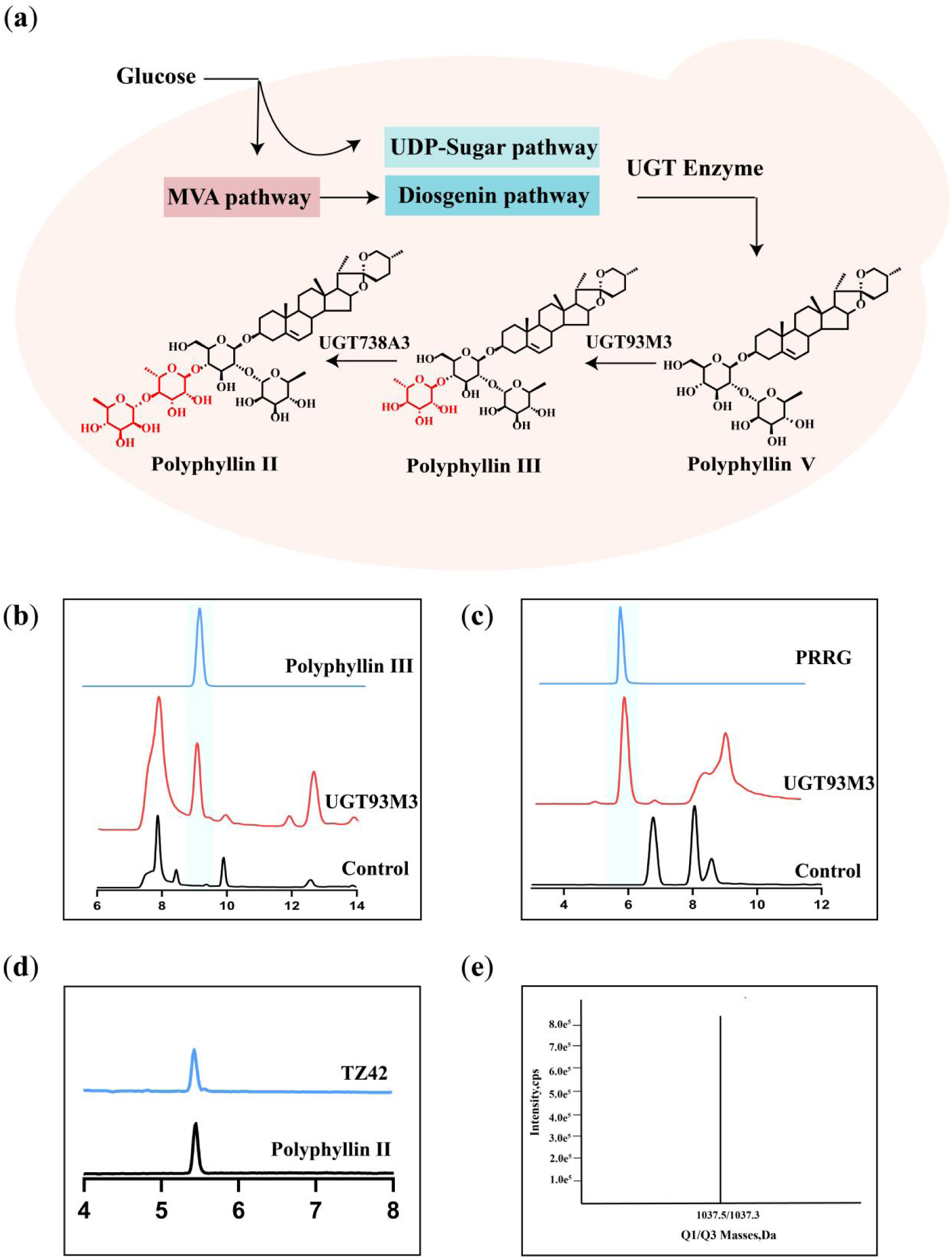
Reconstruction of the polyphyllin II biosynthetic pathway in engineered yeast strain. (a) Coexpression of UGT93M3 and UGT738A3 in yeast strain. (b) HPLC analysis confirms UGT93M3’s role in transforming polyphyllin V into polyphyllin III. (c) HPLC analysis confirms UGT93M3’s role in transforming polyphyllin VI into PRRG . (d) HPLC analysis of TZ42 strain. (e) Mass spectrum of TZ42 strain.

## Discussion

Plants synthesize a wide range of secondary metabolites with pharmacological properties via complex biosynthetic pathways. Nevertheless, the complexity of plant genomes and the limited availability of genomic data have restricted research into the biosynthetic pathways of these metabolites. The medicinal plant *T. tschonoskii* is rich in polyphyllins, which are the primary active components responsible for its therapeutic efficacy on Alzheimer’s disease, and confer anti-aging, anti-tumor, anti-inflammatory, and central nervous system regulatory activities. Recently, a *RhaGT* gene, which is responsible for the formation of tetraglycoside, was identified in *Paris polyphylla*(Zhou *et al*., 2024.). However, the biosynthetic pathway of steroidal saponins mediated by GTs remains to be fully elucidated. Molecular-level research on this species is still limited. Therefore, clarifying the biosynthetic pathway of polyphyllins has become an important research focus. In this study, we sequenced the transcriptome of *T. tschonoskii* and successfully identified a highly specific RhaGT gene for 4’’-OH based on gene annotation information and metabolite content determination..Furthermore, by introducing UGT93M3 and UGT738A3 into yeast strains engineered to produce polyphyllin V, the *in vivo* functionality of UGT738A3 was subsequently validated.

UGT738A3 exhibits relatively high substrate specificity. Most reported GTs have broad substrate promiscuity. For instance, SgUGT94-289-3 adopts a dual-pocket catalytic mechanism where the N-terminal α-helix swings to increase the pocket volume, enabling it to recognize at least five different substrates and demonstrating strong promiscuity (Cui *et al*., 2024). As a tetra-glycosyltransferase for sugar chain extension, UGT738A3 has a predicted pocket volume of only 612.86 Å³, which makes it difficult for branched and linear glycoside compounds to enter the pocket and reach the catalytic center. It also makes it hard for the compounds in the pocket to undergo conformational changes to expose hydroxyl groups at other positions. Therefore, UGT738A3 exhibits a relatively narrow substrate specificity and demonstrates strong selectivity towards its sugar donor. However, to deeply understand the molecular mechanism of the function of the tetra-glycosyltransferase, further crystal structure analysis is required.

In conclusion, this study successfully identifies the key RhaGT gene UGT738A3 in the polyphyllin biosynthetic pathway and its optimized mutant, thereby achieving complete heterologous biosynthesis of polyphyllin II in engineered yeast. This research not only provides essential enzyme components for the heterologous production of polyphyllins but also offers a robust strategy for elucidating the intricate biosynthetic pathways of other plant metabolites.

## Acknowledgements

This work was supported by the Beijing Natural Science Foundation (7232264), the ability establishment of sustainable use for valuable Chinese medicine resources (2060302), the National Natural Science Foundation of China (81974515), the Central Laboratory of Capital Medical University.

## Author contributions

XZ conceived and initiated the study. YY and HW performed the experiments. X Wang conducted the MD simulation.W Huang conducted the protein crystal structure determination.X Liu and H Zhao provided guidance.YY wrote the manuscript.XZ, YH and YT revised the manuscript.

## Supporting Information

Additional Supporting Information may be found online in the Supporting Information section at the end of the article.

**Fig. S1**Transcriptome Sequencing Analysis.

**Fig. S2** Four structural formulas were utilized for content determination.

**Fig. S3** Phylogenetic analysis and construction of the UGT gene family tree.

**Fig. S4** HeatMap depicting the expression profile of RhaGTs in the transcriptome of *T. tschonoskii*.

**Fig. S5** Cloning and protein expression of UGT738A3.

**Fig. S6** Enzymatic property analysis results.

**Fig. S7** Protein purification results.

**Fig. S8** Structural formula of the compound.

**Fig.S9** Homologous sequence comparison between UGT738A3 sequence and rhamnosyltransferase genes.

**Fig.S10** Saturation mutagenesis of residue A158.

**Fig. S11** Functional validation and docking results of the A158T mutant.

**Fig. S12** The catalytic results of protodioscin by UGT738A3^A158T/P101L^ are presented

**Fig. S13** Analysis of polyphyllin V/VI Biosynthetic Pathway.

**Table S1** The reported rhamnosyltransferase gene.

**Table S2** Nucleotide sequences of UGT738A3.

**Table S3** Protein sequences of UGT738A3.

**Table S4** Primers used in this work.

**Table S5** Predicted protein pocket volume and depth.

