## Supplementary material for "Identification and Engineering of UDP-rhamnosyltransferase from *Trillium tschonoskii* for Heterologous Biosynthesis of Polyphyllin II in Engineered Yeast": Fig. S1-Fig. S13;Table S1-Table S5

The following Supporting Information is available for this article:

**Fig. S1**Transcriptome Sequencing Analysis.

**Fig. S2** Four structural formulas were utilized for content determination.

**Fig. S3** Phylogenetic analysis and construction of the UGT gene family tree.

**Fig. S4** HeatMap depicting the expression profile of RhaGTs in the transcriptome of *T. tschonoskii.*

**Fig. S5** Cloning and protein expression of UGT738A3.

**Fig. S6** Enzymatic property analysis results.

**Fig. S7** Protein purification results.

**Fig. S8** Structural formula of the compound.

**Fig.S9** Homologous sequence comparison between UGT738A3 sequence and rhamnosyltransferase genes.

**Fig.S10** Saturation mutagenesis of residue A158.

**Fig. S11** Functional validation and docking results of the A158T mutant.

**Fig. S12** The catalytic results of protodioscin by UGT738A3$^{A158T/P101L}$ are presented

**Fig. S13** Analysis of polyphyllin V/VI Biosynthetic Pathway.

**Table S1** The reported rhamnosyltransferase gene.

**Table S2** Nucleotide sequences of UGT738A3.

**Table S3** Protein sequences of UGT738A3.

**Table S4** Primers used in this work.

**Table S5** Predicted protein pocket volume and depth.

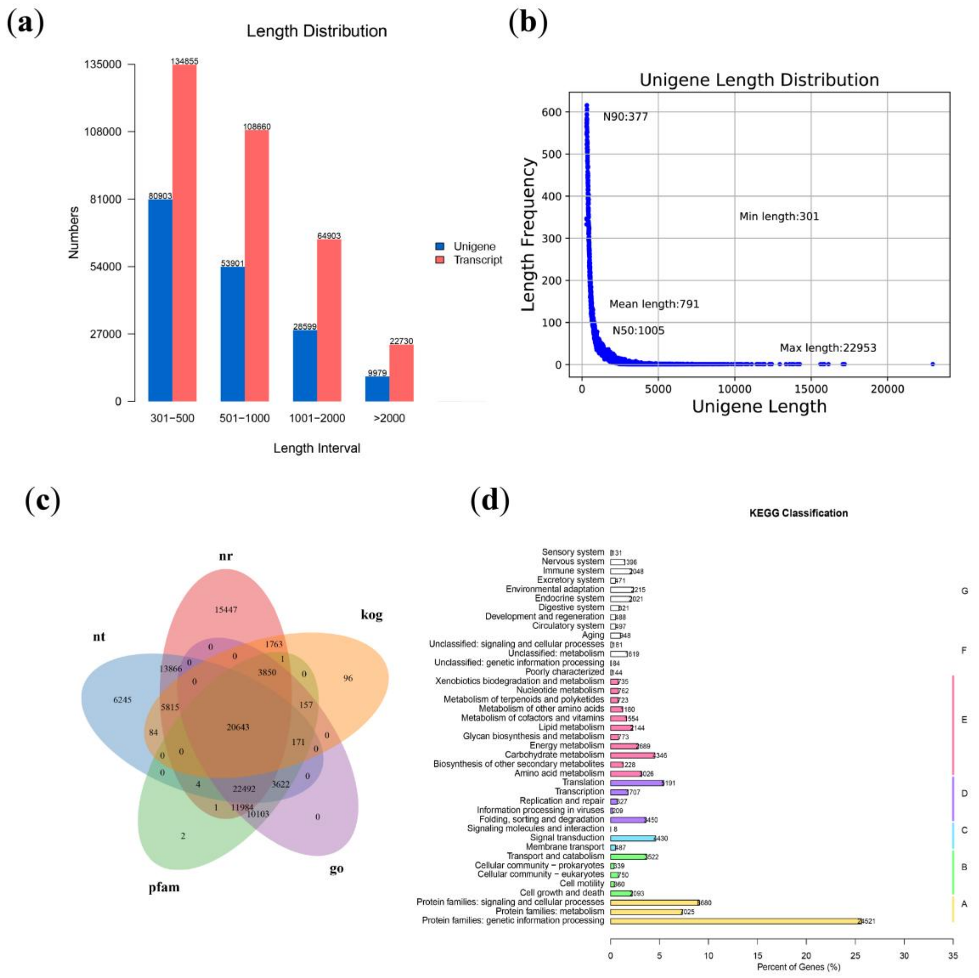


**Fig.S1** Transcriptome Sequencing Analysis.（a）Length distribution of raw reads.（b）Unigene length distribution. （c）Venn diagram for annotation results. （d）KEGG pathway annotation results. Categories include: A, Cellular Processes; B, Environmental Information Processing; C, Genetic Information Processing; D, Metabolism;E,Organismal Systems

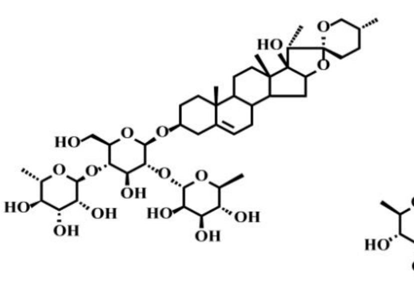

Pennogenin 3-o-beta-chacotrioside

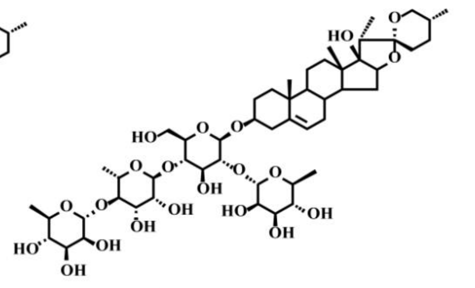

Polyphyllin VII

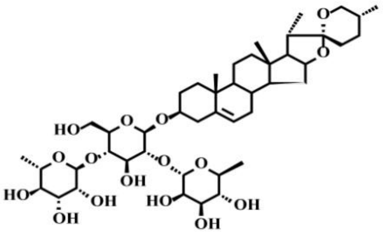

Polyphyllin III

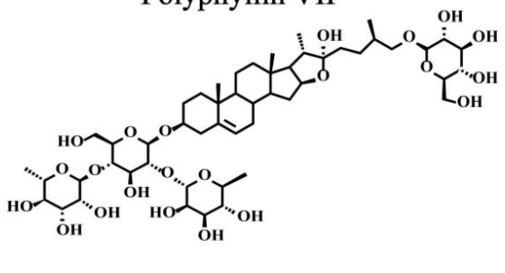

Protodioscin

**Fig. S2** Four structural formulas were utilized for content determination.

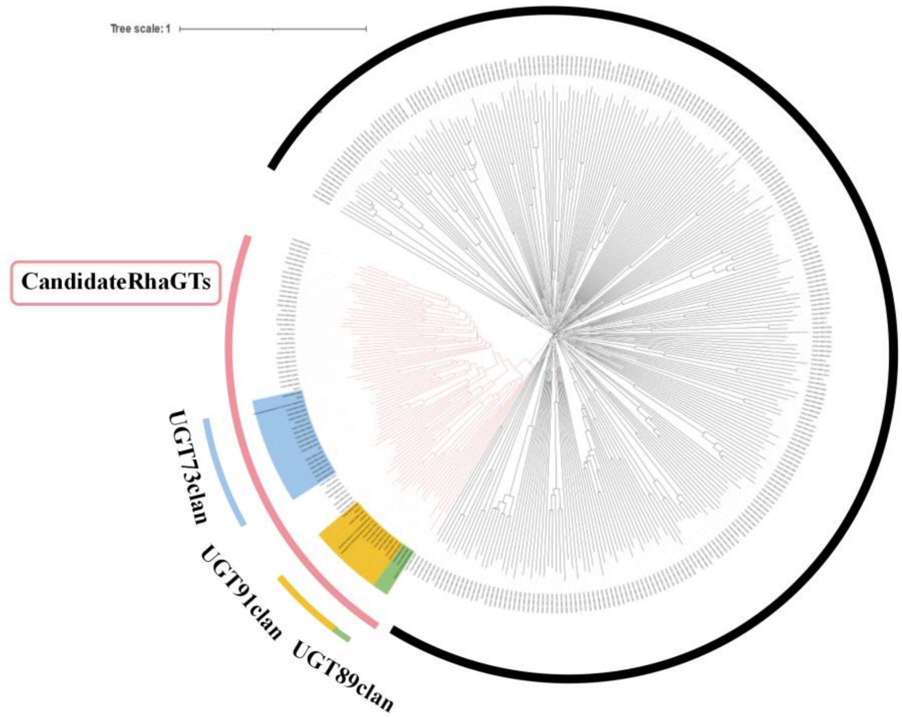


**Fig. S3** Phylogenetic analysis and construction of the UGT gene family tree.

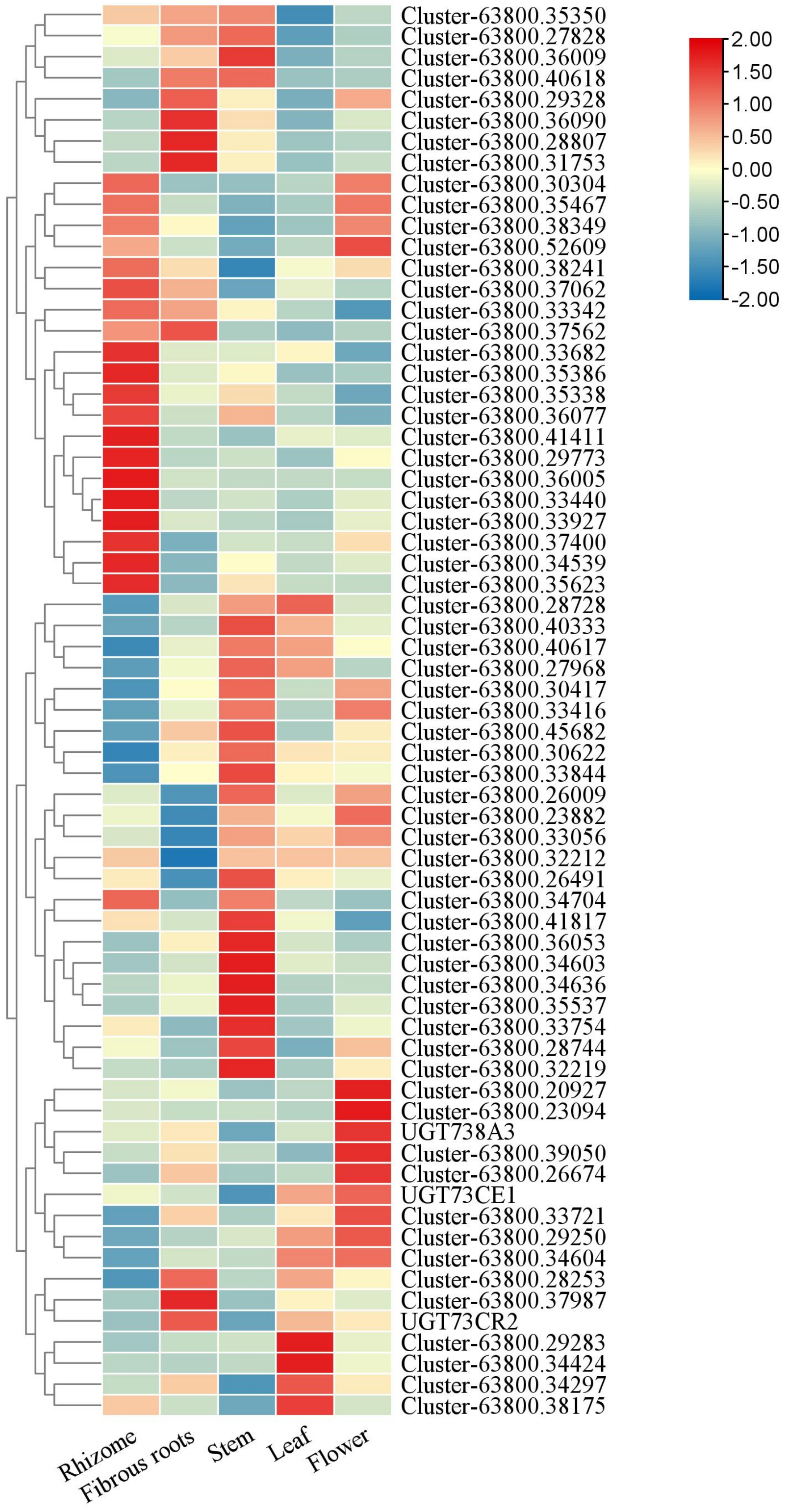


**Fig.S4** HeatMap depicting the expression profile of RhaGTs in the transcriptome of *T. tschonoskii.*

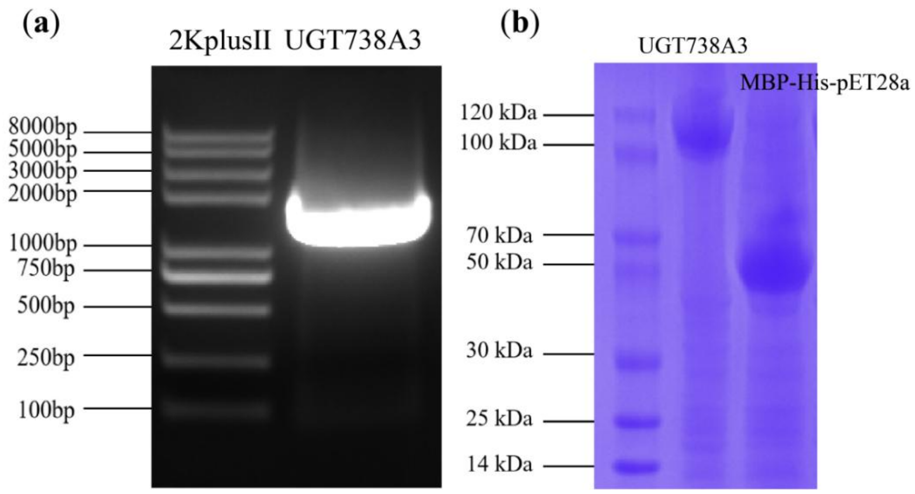


**Fig.S5** Cloning and protein expression of UGT738A3.（a） Amplification of the full-length sequence of UGT738A3.（b）Expression results of the recombinant protein MBP-His-pET28a-UGT738A3.

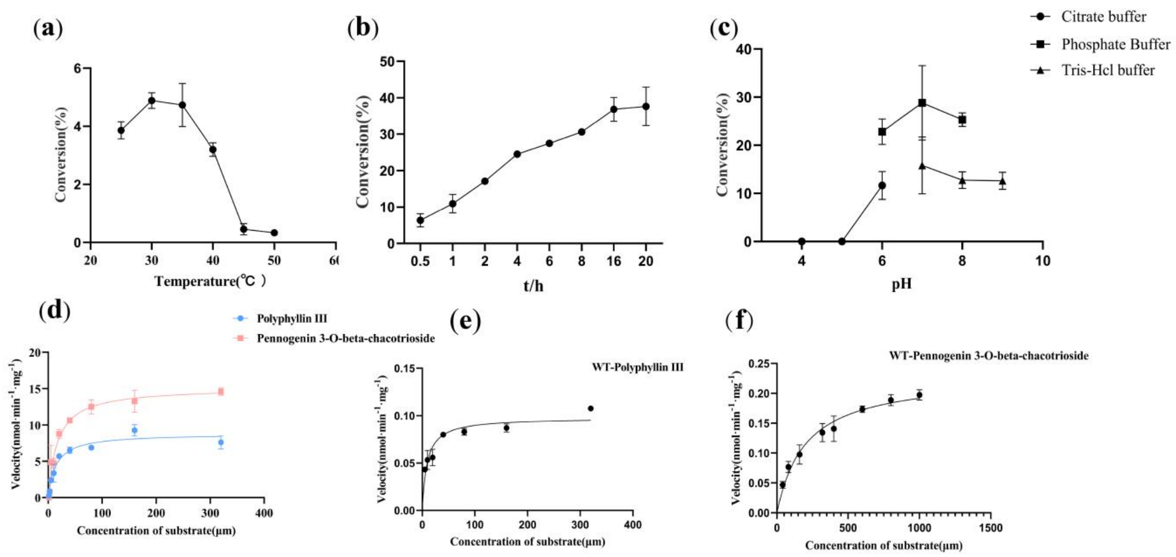


**Fig.S6** Enzymatic property analysis results.（a）The effect of reaction temperature on enzymatic activity.（b）The influence of reaction time on enzymatic activity.（c）The impact of pH on enzymatic activity.（d）Kinetic parameters for UGT738A3$^{A158T/P101L}$ using polyphyllin III and pennogenin 3-O-beta-chacotrioside as substrates, with UDP-Rha serving as the sugar donor.（e）Kinetic parameters for UGT738A3. using polyphyllin III as a substrate and UDP-Rha as the sugar donor.（f）Kinetic parameters for UGT738A3 using pennogenin 3-O-beta-chacotrioside as a substrate and UDP-Rha as the sugar donor.

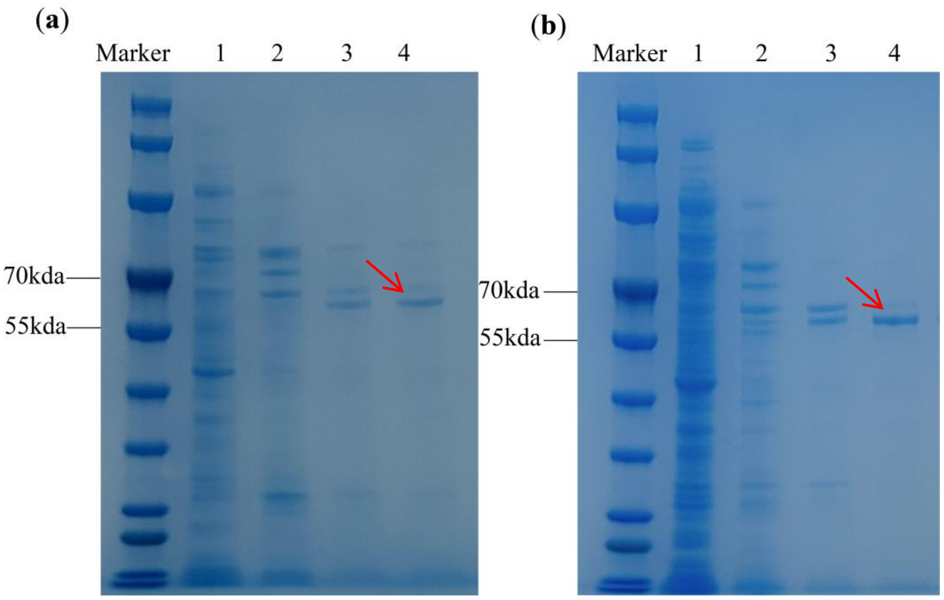


**Fig. S7** Protein purification results. （a） Purification results of the His-pET28a-UGT738A3 protein. （b） Purification results of the His-pET28a-UGT738A3$^{A158T/P101L}$ protein.Lane 1~4 represent elution fractions containing 10mM, 50mM, 100mM, 200mM imidazole, respectively

(a)

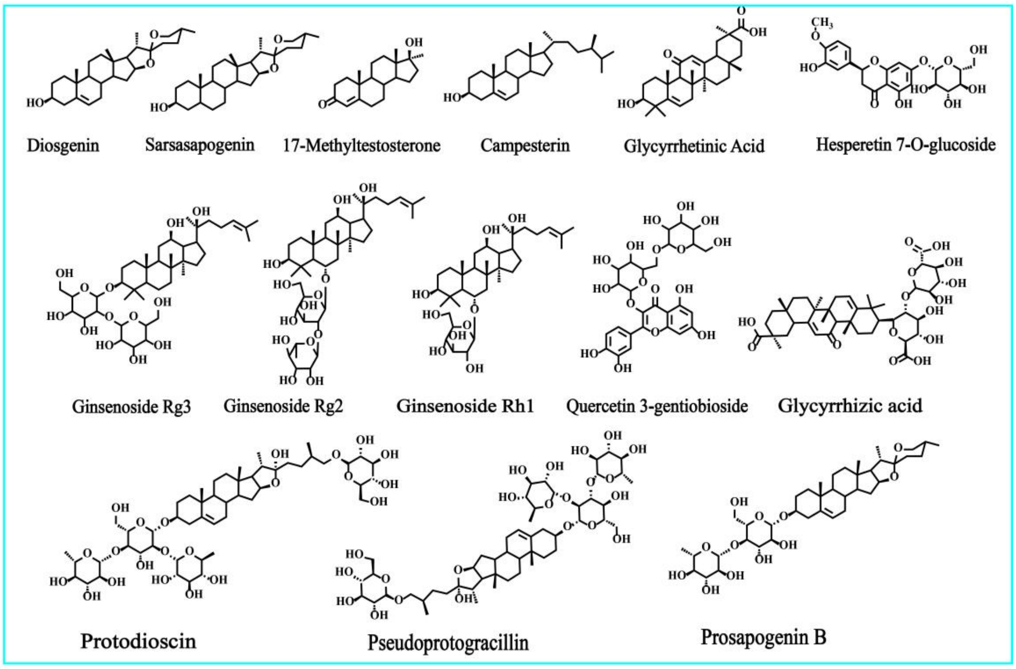


(b)

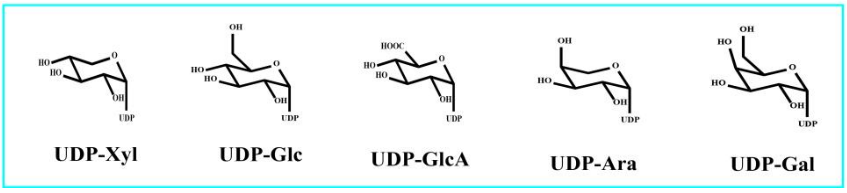


**Fig. S8** Structural formula of the compound.（a）Structural formulae of the chemicals used in the substrate breadth experiments.（b）Five sugar donors employed for glycosylation reactions.

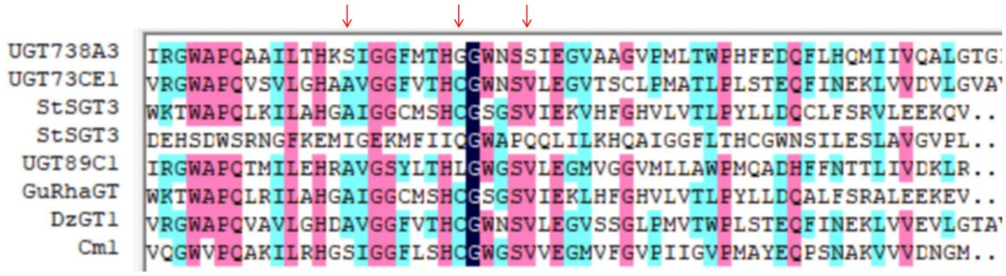


**Fig.S9** Homologous sequence comparison between UGT738A3 sequence and rhamnosyltransferase genes.

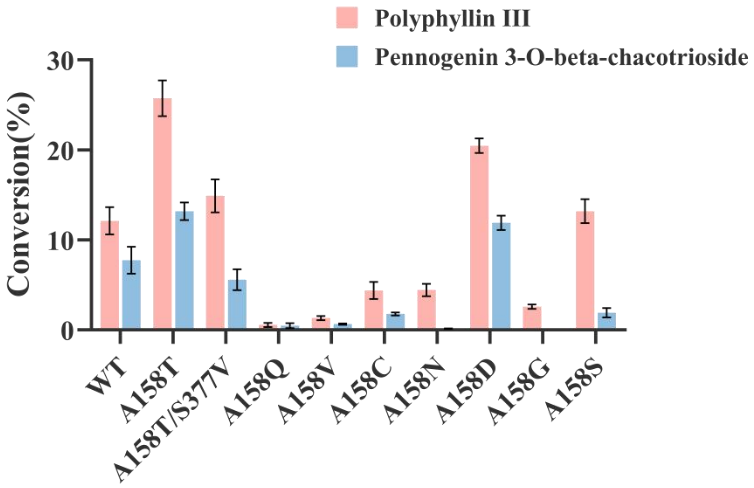


**Fig.S10** Saturation mutagenesis of residue A158.

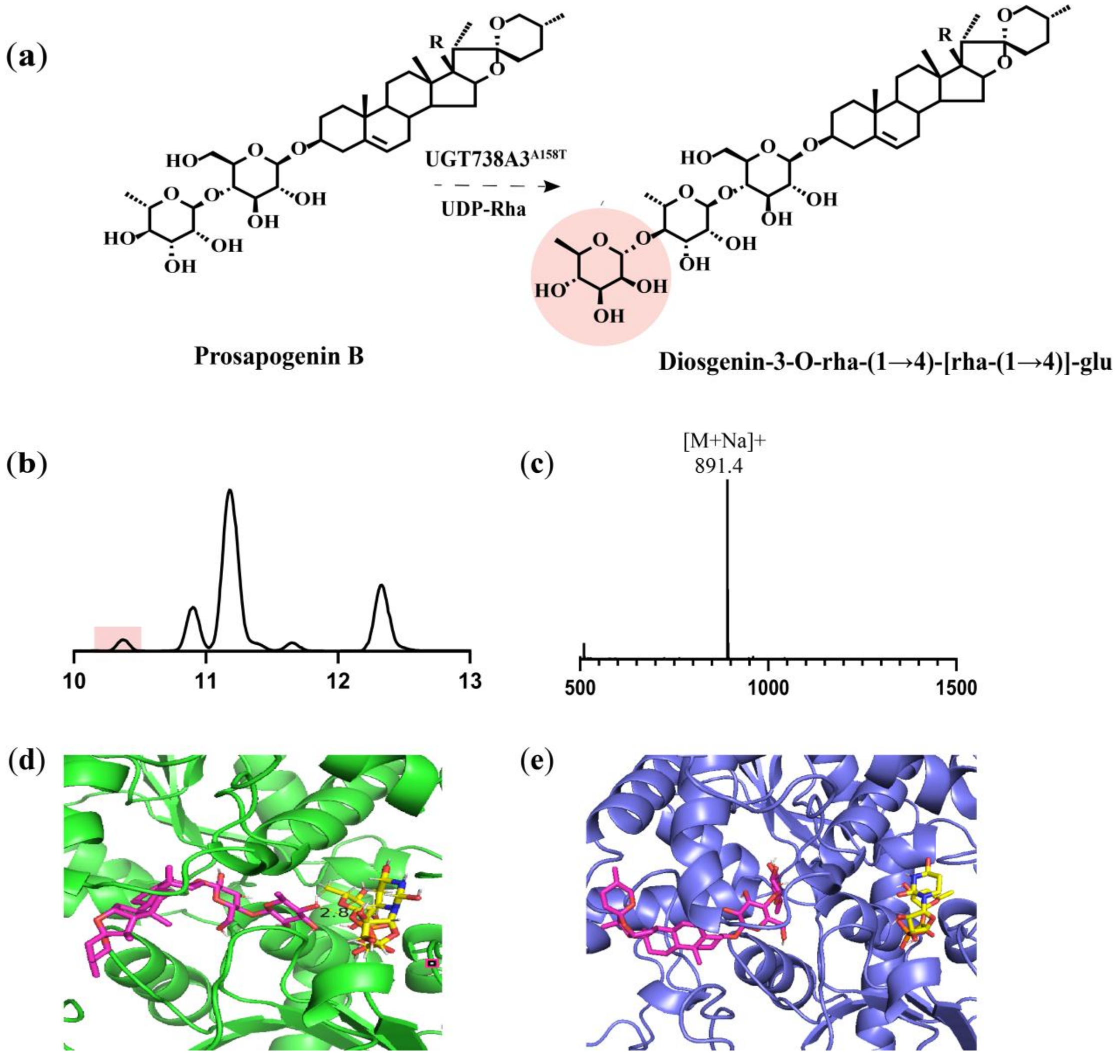


**Fig. S11** Functional validation and docking results of the A158T mutant.（a）The A158T mutant catalyzes the putative products of prosapogenin B.（b）Chromatogram of the product generated from A158T-catalyzed prosapogenin B. (c) Mass spectra of the products derived from A158T-catalyzed prosapogenin B.（d）Docking results between A158T mutant protein and prosapogenin B.（e）Docking results between the WT protein and prosapogenin B.

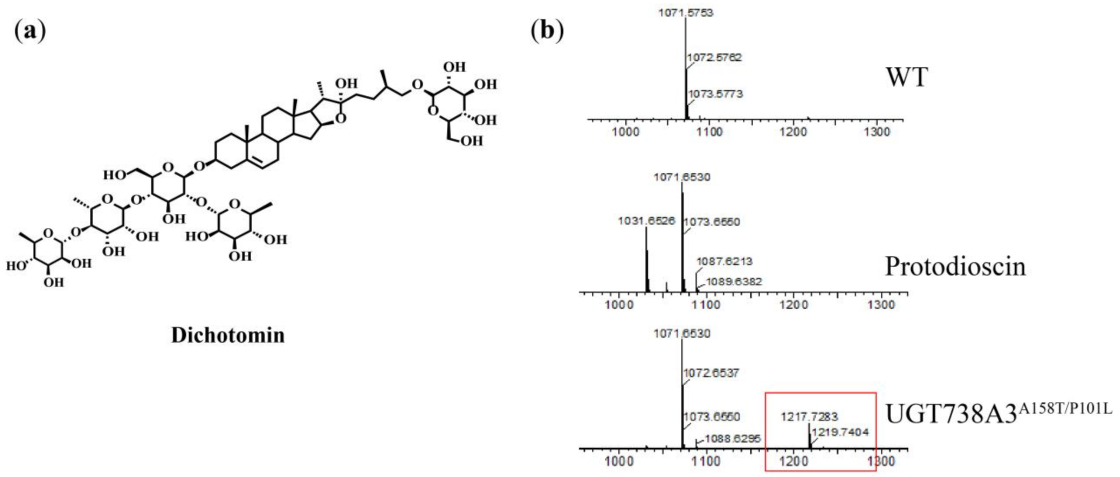


**Fig. S12** The catalytic results of protodioscin by UGT738A3$^{A158T/P101L}$ are presented. （a）Schematic representation of the product structure.（b）Mass spectrometry analysis of the detected product .

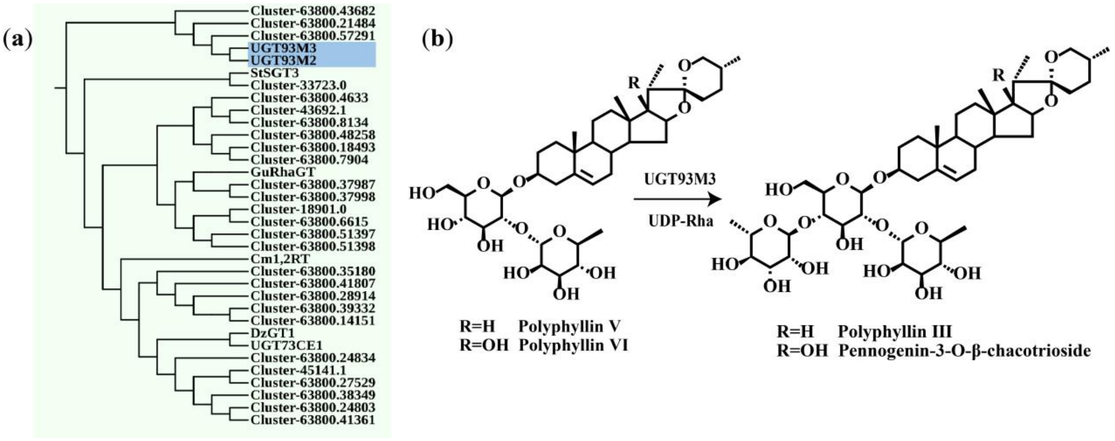


**Fig. S13** Analysis of polyphyllin V/VI Biosynthetic Pathway.（a）RhaGTs closely related to UGT93M2/3.（b）UGT93M3 catalyzes the conversion of polyphyllin V/VI into polyphyllin III and pennogenin 3-O-beta-chacotrioside.

**Table S1** The reported rhamnosyltransferase gene

| Name | Species | Function | Gene ID |
|---|---|---|---|
| UGT89C1 | Arabidopsis thaliana | 7-O-rhamnosyltransferase | 837109 |
| DzGT1 | Dioscorea zingiberensis | rhamnosyltransferase | QZL13805 |
| StSGT3 | Solanum tuberosum | solanidine UDP-glucose glucosyltransferase 1 | P93789 |
| Cm1,2RT | Citrus maxima | flavanone 7-O-glucoside 2''-O-β-L-rhamnosyltransferase | Q8GVE3 |
| GuRhaGT | Glycyrrhiza uralensi | triterpenoid saponin 2''-O-rhamnosyltransferase | UOH28391.1 |
| UGT73CE1 | Paris polyphylla | UDP-rhamnosyltransferase | WIL59760.1 |
| UGT93M3 | S. melongena | UDP-rhamnosyltransferase | |
| UGT93M2 | S. melongena | UDP-rhamnosyltransferase | |

**Table S2 Nucleotide sequences of UGT738A3.**

| Gene Name | Nucleotide sequence |
|---|---|
| UGT738A3 | ATGGGGAGCGCTCCTCCTGTTGAGAGCAGCCAGAAGCACTTC<br>GTGCTGGTCCCGTGGCTGTCCCACGGGCACGTGATCCC<br>GATGATGGACATGGCCCGCCTCCTCGCCGAGCGCGGCGGCA<br>TCCACGTGACCGTGGCCATCTCCCCCGTGGGGGCGGAGC<br>GCATCCGGAGCTGCTTCATCGAGCCCGTCACCGCCGCGAAGC<br>TCCCCATCTCCTTCCTCGACCTCCCCTTCCCCTGCGCC<br>GAAGCTGGGCTCCCGGACGGCATCGAGACCATCGAGCAAAT<br>CCAGGACCCGTCCCTGTTCCCCAAGATGCACGTCGCCGC<br>TGGCCTCCTCAGCAAACCCCTCGAGTCCAAGCTCCGAGAGCT<br>CCCCCGCAAGCCCTCCGTCATCCTCGCCGACCTCTACC<br>ACCCGTGGGCGCGGGAAGTCGCGGCCGACCTCGGCGTCCCG<br>CTGCTGCTCTACTACGTGTTCCCGTGCTTCGCCATCCTC<br>GTCTACCGCAGTCTGAGACAGCATGGTGTCTACGATGACGGC<br>GCGGCGGACGCGAGCCGGATGTTCCCGGTGCCTGACGC<br>GCCGGAGTACATGGTCAGCCGGGCGCAGGCGCCGGGGACGT<br>TCGACAGGCCCGGGTGGGAGTGGCTTCGAGAGGAAGCTA<br>TTGCGGCTGAGTCCGCCGCCGCCGGGGTTATTTTCACAGCT<br>TCGACCAGCTCGAGCCCAATTTCCTCCCCAAGTTCCAG<br>GAGATCATGGGGGGCCTGAAGACGTGGGCCATCGGCCCGCT<br>GTCCCTCAGCCACAAGGACGTGCTGGCCGAGCGCGGGAG<br>CGCAAATGAGGTCGCCGCCGACCGCTGCCTCACCTGGCTCGA<br>CGCCAACGCTCCCGCCTCCGTCATCTACGTCTGCTTCG<br>GCACCAACACATACTGGACCCCCCAGCAGATCATCGAGGTC<br>GGGTCCGGGATCGAGAGCTCGGGCCACCCATTCATCTGG<br>GTGCTGAAGAAGCGGGAGCTGACGCCGGAGGTGGAGGAGTT<br>CCTGTCGGGAGGGTTCGAGGAGCGGACGCGGGACCGAGG<br>CCTGCTCATCAGGGGCTGGGCCCCTCAGGCGGCCATACTGAC<br>CCATAAGTCGATCGGGGGATTCATGACACATGGCGGGT<br>GGAACTCGTCGATCGAAGGGGTGGCGGCCGGGGTGCCAATG<br>CTGACGTGGCCGCACTTCGAGGACCAGTTCTTGCACCAG<br>ATGATCATCGTGCAGGCGCTGGGGACGGGGATCGGGGTCGG<br>GGTGCGGGCGCAGGAGGACTACATCGCGCAGGTGATGGA<br>CACCATCAAGCGGGAGCAGGTGGAGAAGGCCGTGAGGGAGC<br>TGATGGGACGAGGGGAGGAAGCCGACGCGAGGAGGAGAA<br>AGGCCAAGGAGTACGGGGAGAAGGCGAGGAGGGCAATGGA<br>GGTCGGGGGGTCGTCGTATGTGAATTTGACCGAAGTGATC<br>GACTCGATTCCAGCTGTTGTCGCCACCGAGAATGGTGGTGGT<br>GACTAA |

**Table S3 Protein sequence sequences of UGT738A3.**

| Gene Name | Protein sequence |
|---|---|
| UGT738A3 | MGSAPPVESSQKHFVLVPWLSHGHVIPMMDMARLLAERGGIH<br>VTVAISPVGAERIRSCFIEPVTAAKLPISFLDLPFPCAEAGLPDGIE<br>TIEQIQDPSLFPKMHVAAGLLSKPLESKLRELPRKPSVILADLYH<br>PWAREVAADLGVPLLLYYVFPCFAILVYRSLRQHGVYDDGAA<br>DASRMFPVPDAPEYMVSRAQAPGTFDRPGWEWLREEAIAAESA<br>AAGVIFHSFDQLEPNFLPKFQEIMGGLKTWAIGPLSLSHKDVLA<br>ERGSANEVAADRCLTWLDANAPASVIYVCFGTNTYWTPQQIIE<br>VGSGIESSGHPFIWVLKKRELTPEVEEFLSGGFEERTRDRGLLIR<br>GWAPQAAILTHKSIGGFMTHGGWNSSIEGVAAGVPMLTWPHFE<br>DQFLHQMIIVQALGTGIGVGVRAQEDYIAQVMDTIKREQVEKA<br>VRELMGRGEEADARRRKAKEYGEKARRAMEVGGSSYVNLTE<br>VIDSIPAVVATENGGGD |

**Table S4** Primers used in this work.

| Primer name | Sequence（5'-3'） |
|---|---|
| UGT738A3-F | AAAAAGGACAAAAAACTATTCACCC |
| UGT738A3-R | CACACCAAAAAGATGAATGAAGAGA |
| S376A-F | CATGGCGGGTGGAACGCGTCGATCGA |
| S376A-R | CGTTCCACCCGCCATGTGTCATGAAT |
| T370A-F | ATCGGGGGATTCATGGCACATGGCGG |
| T370A-R | CCATGAATCCCCCGATCGACTTATGG |
| R33A-F | ATGATGGACATGGCCGCCCTCCTCGCC |
| R33A-R | GCGGCCATGTCCATCATCGGGATCACG |
| H22A-F | GTCCCGTGGCTGTCCGCCGGGCACGTG |
| H22A-R | GCGGACAGCCACGGGACCAGCACGAAG |
| P355A-F | ATCAGGGGCTGGGCCGCTCAGGCGGC |
| P355A-R | CGGCCCAGCCCCTGATGAGCAGGCCT |
| P27A-F | CACGGGCACGTGATCGCGATGATGGA |
| P27A-R | CGATCACGTGCCCGTGGGACAGCCAC |
| I26A-F | TCCCACGGGCACGTGGCCCCGATGATG |
| I26A-R | CGCACGTGCCCGTGGGACAGCCACGGG |
| E379A-F | GGAACTCGTCGATCGCAGGGGTGGCG |
| E379A-R | GCGATCGACGAGTTCCACCCGCCATG |
| R54A-F | CCCGTGGGGGCGGAGGCCATCCGGAGC |
| R54A-R | GCCTCCGCCCCCACGGGGGAGATGGCC |
| V50A-F | TGGCCATCTCCCCCGCGGGGGCGGAG |
| V50A-R | GCGGGGGAGATGGCCACGGTCACGTG |
| E53A-F | CCCCCGTGGGGGCGGCGCATCCGG |
| E53A-R | GCCGCCCCCACGGGGGAGATGGCCAC |
| S255A-F | GCCATCGGCCCGCTGGCCCTCAGCCA |
| S255A-R | CCAGCGGGCCGATGGCCCACGTCTTC |
| G23A-F | CGTGGCTGTCCCACGCGCACGTGATC |
| G23A-R | GCGTGGGACAGCCACGGGACCAGCAC |
| A158T-F | GTGTTCCCGTGCTTCACCATCCTCGT |
| A158T-R | TGAAGCACGGGAACACGTAGTAGAGC |
| I91A-F | GACGGCATCGAGACCGCCGAGCAAATC |
| I91A-R | GCGGTCTCGATGCCGTCCGGGAGCCCA |

| | |
|---|---|
| F100A-F | CAGGACCCGTCCCTGGCCCCCAAGATG |
| F100A-R | GCCAGGGACGGGTCCTGGATTTGCTCG |
| H104A-F | CTGTTCCCCAAGATGGCCGTCGCCGCT |
| H104A-R | GCCATCTTGGGGAACAGGGACGGGTCC |
| L132A-F | GTCATCCTCGCCGACGCCTACCACCCG |
| L132A-R | GCGTCGGCGAGGATGACGGAGGGCTTG |
| Y152A-F | CCGCTGCTGCTCTACGCCGTGTTCCCG |
| Y152A-R | GCGTAGAGCAGCAGCGGGACGCCGAGG |
| F154A-F | CTGCTCTACTACGTGGCCCCGTGCTTC |
| F154A-R | GCCACGTAGTAGAGCAGCAGCGGGACG |
| I159A-F | TTCCCGTGCTTCGCCGCCCTCGTCTAC |
| I159A-R | GCGGCGAAGCACGGGAACACGTAGTAG |
| P198A-F | AGCCGGGCGCAGGCGGCGGGGACGTT |
| P198A-R | CCGCCTGCGCCCGGCTGACCATGTAC |
| T200A-F | GCGCAGGCGCCGGGGGCGTTCGACAG |
| T200A-R | CCCCCGGCGCCTGCGCCCGGCTGACC |
| F201A-F | CAGGCGCCGGGGACGGCCGACAGGCCC |
| F201A-R | GCCGTCCCCGGCGCCTGCGCCCGGCTG |
| R203A-F | CCGGGGACGTTCGACGCGCCCGGGTGG |
| R203A-R | GCGTCGAACGTCCCCGGCGCCTGCGCC |
| L209A-F | CCCGGGTGGGAGTGGGCTCGAGAGGAA |
| L209A-R | GCCCACTCCCACCCGGGCCTGTCGAAC |
| N296A-F | GTCTGCTTCGGCACCGCCACATACTGG |
| N296A-R | GCGGTGCCGAAGCAGACGTAGATGACG |
| F393A-F | CTGACGTGGCCGCACGCCGAGGACCAG |
| F393A-R | GCGTGCGGCCACGTCAGCATTGGCACC |
| Y421A-F | CGGGCGCAGGAGGACGCCATCGCGCAG |
| Y421A-R | GCGTCCTCCTGCGCCCGCACCCCGACC |
| A158R-F | GTGTTCCCGTGCTTCCGCATCCTCGTC |
| A158R-R | CGGAAGCACGGGAACACGTAGTAGAGC |
| A158N-F | GTGTTCCCGTGCTTCAACATCCTCGTC |
| A158N-R | TTGAAGCACGGGAACACGTAGTAGAGC |
| A158D-F | GTGTTCCCGTGCTTCGACATCCTCGTC |

| | |
|---|---|
| A158D-R | TCGAAGCACGGGAACACGTAGTAGAGC |
| A158C-F | GTGTTCCCGTGCTTCTGCATCCTCGTC |
| A158C-R | CAGAAGCACGGGAACACGTAGTAGAGC |
| A158Q-F | GTGTTCCCGTGCTTCCAGATCCTCGTCT |
| A158Q-R | CTGGAAGCACGGGAACACGTAGTAGAGC |
| A158E-F | GTGTTCCCGTGCTTCGAGATCCTCGTCT |
| A158E-R | CTCGAAGCACGGGAACACGTAGTAGAGC |
| A158G-F | TGTTCCCGTGCTTCGGCATCCTCGTC |
| A158G-R | CCGAAGCACGGGAACACGTAGTAGAG |
| A158H-F | GTGTTCCCGTGCTTCCACATCCTCGTC |
| A158H-R | TGGAAGCACGGGAACACGTAGTAGAGC |
| A158I-F | GTGTTCCCGTGCTTCATCATCCTCGTC |
| A158I-R | ATGAAGCACGGGAACACGTAGTAGAGC |
| A158L-F | GTGTTCCCGTGCTTCCTCATCCTCGTC |
| A158L-R | AGGAAGCACGGGAACACGTAGTAGAGC |
| A158K-F | GTGTTCCCGTGCTTCAAGATCCTCGTCT |
| A158K-R | CTTGAAGCACGGGAACACGTAGTAGAGC |
| A158M-F | GTGTTCCCGTGCTTCATGATCCTCGTCT |
| A158M-R | CATGAAGCACGGGAACACGTAGTAGAGC |
| A158F-F | GTGTTCCCGTGCTTCTTCATCCTCGTC |
| A158F-R | AAGAAGCACGGGAACACGTAGTAGAGC |
| A158P-F | GTGTTCCCGTGCTTCCCCATCCTCGTC |
| A158P-R | GGGAAGCACGGGAACACGTAGTAGAGC |
| A158S-F | GTGTTCCCGTGCTTCTCCATCCTCGTC |
| A158S-R | GAGAAGCACGGGAACACGTAGTAGAGC |
| A158W-F | GTGTTCCCGTGCTTCTGGATCCTCGTCT |
| A158W-R | CCAGAAGCACGGGAACACGTAGTAGAGC |
| A158Y-F | GTGTTCCCGTGCTTCTACATCCTCGTC |
| A158Y-R | TAGAAGCACGGGAACACGTAGTAGAGC |
| A158V-F | GTGTTCCCGTGCTTCGTCATCCTCGTC |
| A158V-R | ACGAAGCACGGGAACACGTAGTAGAGC |
| W19A-F | TTCGTGCTGGTCCCGGCGCTGTCCCAC |
| W19A-R | GCCGGGACCAGCACGAAGTGCTTCTGG |

| | |
|---|---|
| W19L-F | TCGTGCTGGTCCCGTTGCTGTCCCAC |
| W19L-R | AACGGGACCAGCACGAAGTGCTTCTG |
| W19F-F | TCGTGCTGGTCCCGTTCCTGTCCCACG |
| W19F-R | GAACGGGACCAGCACGAAGTGCTTCTG |
| F201A-F | CAGGCGCCGGGGACGGCCGACAGGCCC |
| F201A-R | GCCGTCCCCGGCGCCTGCGCCCGGCTG |
| F201L-F | CAGGCGCCGGGGACGCTCGACAGGCC |
| F201L-R | GCGTCCCCGGCGCCTGCGCCCGGCTG |
| P101A-F | GACCCGTCCCTGTTCGCCAAGATGCA |
| P101A-R | CGAACAGGGACGGGTCCTGGATTTGC |
| P101L-F | ACCCGTCCCTGTTCCTCAAGATGCAC |
| P101L-R | AGGAACAGGGACGGGTCCTGGATTTG |
| P101F-F | GACCCGTCCCTGTTCTTCAAGATGCAC |
| P101F-R | AAGAACAGGGACGGGTCCTGGATTTGC |
| Y133A-F | ATCCTCGCCGACCTCGCCCACCCGTGG |
| Y133A-R | GCGAGGTCGGCGAGGATGACGGAGGGC |
| Y133L-F | TCCTCGCCGACCTCTTGCACCCGTGGG |
| Y133L-R | CAAGAGGTCGGCGAGGATGACGGAGGG |
| Y133F-F | TCCTCGCCGACCTCTTCCACCCGTGG |
| Y133F-R | AAGAGGTCGGCGAGGATGACGGAGGG |
| W206A-F | TTCGACAGGCCCGGGGCGGAGTGGCTT |
| W206A-R | GCCCCGGGCCTGTCGAACGTCCCCGGC |
| W206L-F | TCGACAGGCCCGGGTTGGAGTGGCTT |
| W206L-R | AACCCGGGCCTGTCGAACGTCCCCGG |
| W206F-F | TCGACAGGCCCGGGTTCGAGTGGCTTC |
| W206F-R | GAACCCGGGCCTGTCGAACGTCCCCGG |
| E92A-F | GCATCGAGACCATCGCGCAAATCCAG |
| E92A-R | GCGATGGTCTCGATGCCGTCCGGGAG |
| E92L-F | GGCATCGAGACCATCCTGCAAATCCAG |
| E92L-R | AGGATGGTCTCGATGCCGTCCGGGAGC |
| E92F-F | GGCATCGAGACCATCTTCCAAATCCAGG |
| E92F-R | GAAGATGGTCTCGATGCCGTCCGGGAGC |

**Table S5** Predicted protein pocket volume and depth

| Name | Volume（Å³) | Depth(Å) |
|---|---|---|
| UGT738A3 | 612.86 | 25.62 |
| UGT738A3$^{A158T/P101L}$ | 740.86 | 25.62 |
